# The Velvet transcription factor PnVeA regulates necrotrophic effectors and secondary metabolism in the wheat pathogen *Parastagonospora nodorum*

**DOI:** 10.1101/2023.11.13.566949

**Authors:** Shota Morikawa, Callum Verdonk, Evan John, Leon Lenzo, Nicolau Sbaraini, Chala Turo, Hang Li, David Jiang, Yit-Heng Chooi, Kar-Chun Tan

**Author notes:** Corresponding author: K.-C. Tan.

## Abstract

The fungus *Parastagonospora nodorum* causes septoria nodorum blotch on wheat. The role of the fungal Velvet-family transcription factor VeA in *P. nodorum* development and virulence was investigated here. Deletion of the *P. nodorum VeA* ortholog, *PnVeA*, resulted in growth abnormalities including pigmentation, abolished asexual sporulation and highly reduced virulence on wheat. Comparative RNA-Seq and RT-PCR analyses revealed that the deletion of *PnVeA* also decoupled the expression of major necrotrophic effector genes. In addition, the deletion of *PnVeA* resulted in an up-regulation of four predicted secondary metabolite (SM) gene clusters. Using liquid-chromatography mass-spectrometry, it was observed that one of the SM gene clusters led to an accumulation of the mycotoxin alternariol. PnVeA is essential for asexual sporulation, full virulence, secondary metabolism and necrotrophic effector regulation.

## Introduction

Plant pathogenic microorganisms cause severe economic damage to the agricultural sector threatening food security (Savary et al. 2019). *Parastagonospora nodorum* is a pathogenic fungus that causes Septoria nodorum blotch (SNB) of wheat and is considered a model necrotrophic fungal plant pathogen (Quaedvlieg et al. 2013; Solomon et al. 2006). The use of fungicides and the breeding of resistant cultivars have been successful in managing the spread of SNB (Downie et al. 2020). However, continued yield losses caused by SNB prompt the investigation for the discovery of novel control methods (Downie et al. 2020).

*P. nodorum* is a necrotroph which infects susceptible wheat cultivars by causing necrosis of the plant tissue which it subsequently colonises (Oliver et al. 2012). *P. nodorum* spores germinate on the surface of wheat leaves (Oliver et al. 2012). During infection, the fungus produces proteinaceous necrotrophic effectors (NE) that interact with wheat dominant susceptibility receptors, in an inverse gene-for-gene manner, leading to host tissue necrosis (Oliver et al. 2012). In most instances, the interaction between the NE and the product of its corresponding susceptibility gene is not a direct protein–protein interaction, which indicates that these effectors induce cell death through other signalling partners (Faris and Friesen 2020). The interaction between NEs and their corresponding susceptibility genes has been conceptualised as effector-triggered susceptibility and *P. nodorum*–wheat has been used as the model pathosystem to study this immune pathway in plants (Faris and Friesen 2020). The infection cycle is completed by the production of asexual pycnidiospores and sexual ascospores which spread through rain splashes and wind, respectively, to infect other susceptible host plants (Solomon et al. 2006). For crop protection purposes, the removal of dominant susceptibility genes from wheats improves disease resistance (Downie et al. 2020).

Whilst transcription factors (TFs) are ubiquitous across all domains of life (Charoensawan et al. 2010), the structural family Velvet is exclusive to the fungal kingdom (Bayram and Braus 2012). Velvet TFs were first identified and characterised as important developmental regulators through pioneering studies in *Aspergillus* spp. (Calvo et al. 2016; Käfer 1965). While all Velvet TFs share a conserved domain, four distinct classes have been described: Velvet A (VeA), Velvet-like B (VelB), Velvet-like C (VelC) and Viability of Spores A (VosA). Furthermore, interactions between the Velvet TFs have been reported, indicating that complex oligomeric regulatory units can form (Ahmed et al. 2014). The Velvet complex can also interact with other cofactors, such as the methyltransferase LaeA, and has been associated with several regulatory functions including vegetative growth and development, stress tolerance, reproduction and secondary metabolite (SM) biosynthesis (Calvo et al. 2016). Orthologs of the canonical Velvet regulator VeA have been associated with virulence in both biotrophic and necrotrophic plant pathogens (John et al. 2021).

A number of *P. nodorum* TFs have been found to regulate the NE genes *SnToxA* and *SnTox3* but little is known about the transcriptional regulation of the other two major NE genes *SnTox1* and *SnTox267* (John et al. 2021; Richards et al. 2022; Tan and Oliver 2017). Previously, *P. nodorum* TFs such as PhmR and ElcR have been shown to regulate the production of phytotoxic SMs and be required for normal phytopathogenicity (Chooi et al. 2017; Li et al. 2020).

Prior to this study, the role of the Velvet pathway of *P. nodorum* in NE and secondary metabolite regulation and host infection was not known. Therefore, this study aimed to identify and characterise the Velvet VeA ortholog in *P. nodorum* (PnVeA), and to investigate the genes regulated by this putative TF. PnVeA is essential for asexual reproduction, vegetative development and full phytopathogenicity. Furthermore, several NEs, as well as the mycotoxin alternariol are subjected to VeA regulatory control.

## Results

### Identification of Velvet TFs in *P. nodorum* and the deletion of *PnVeA* in *P. nodorum*

Four coding sequences were identified in the *P. nodorum* reference (Bertazonni et el., 2021) which harboured a putative velvet domain (InterPro ID: IPR037525) (**Fig. 1A**). A phylogenetic analysis incorporating characterised Velvet TFs from *A. nidulans*, *A. flavus*, *Botrytis cinerea* and *Magnaporthe oryzae* revealed four clades corresponding to the four distinct Velvet ortholog classes VeA, VelB, VelC and VosA (**Fig. 1B**). As such, the putative *P. nodorum* orthologues were designated PnVeA (SNOG_01807), PnVelB (SNOG_03679), PnVelC (SNOG_12917) and PnVosA (SNOG_06311). PnVeA was selected for further investigation and subsequent functional characterisation, given that it is highly expressed during *P. nodorum* infection, from necrotrophy to sporulation based on a microarray analysis (Ipcho et al. 2012), which aligns with the prominent role of VeA orthologs in other fungi, especially during host-pathogen interaction (Calvo et al. 2016; John et al., 2021).

**Figure 1:**
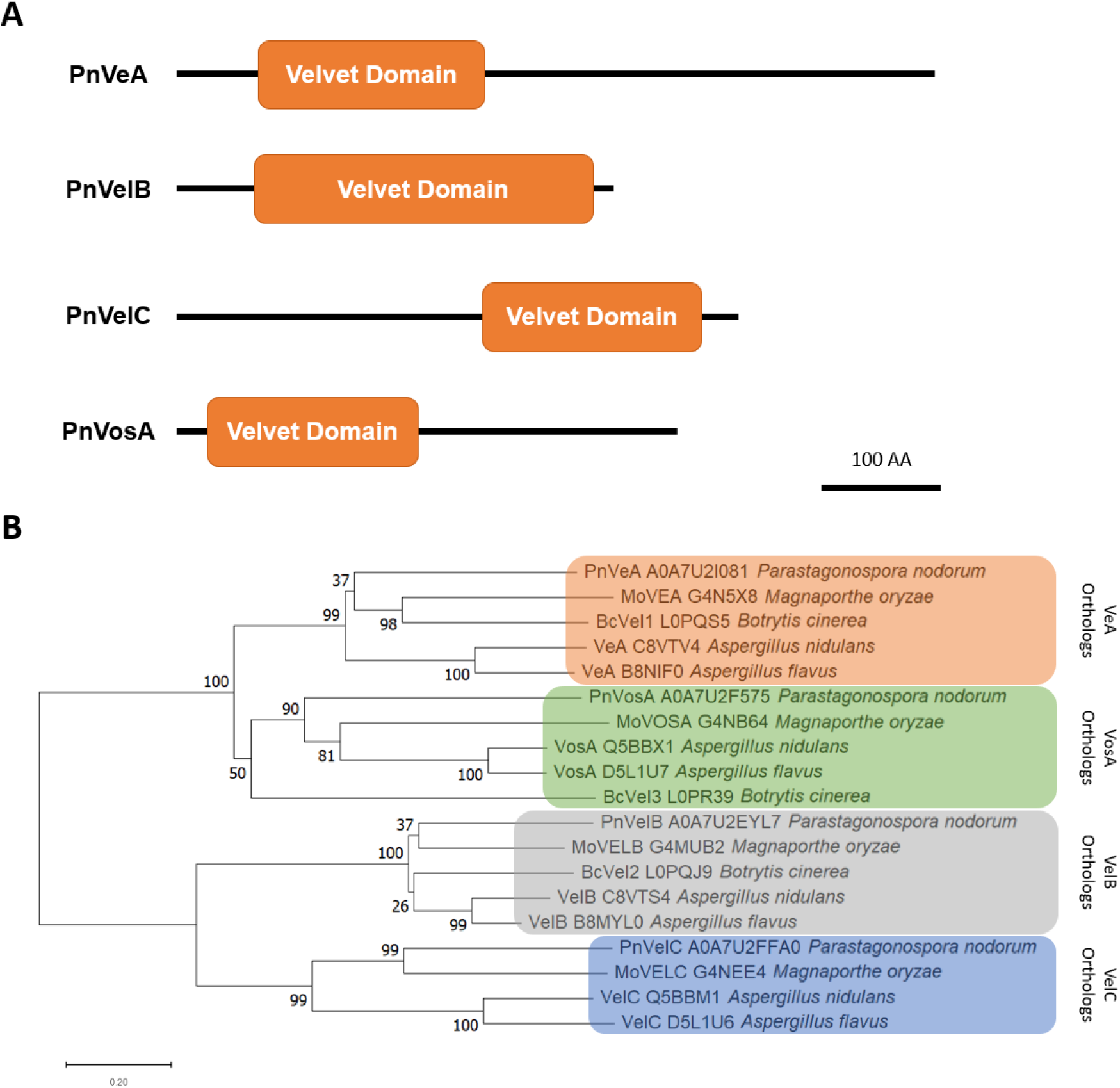
Protein sequence analysis of Velvet transcription factors. **(A)** Domain architecture of putative Velvet-domain (IPR037525) proteins in *Parastagonospora nodorum* showing the location of the Velvet domain in each ortholog. **(B)** A neighbour-joining phylogenetic tree showing characterised Velvet-domain transcription factors from four different species compared to putative orthologues in *P. nodorum*. Orange, green, grey, and blue boxes indicate the clustered VeA, VosA, VelB and VelC orthologs respectively. UniProt IDs are indicated right of protein names. Branch lengths denote the dissimilarities in amino acid sequences between proteins. Bootstrap values are indicated at node branches.

### PnVeA is required for vegetative development, asexual sporulation and full virulence

Two independent *P. nodorum* mutant strains lacking a functional *PnVeA* (SNOG_01807, NCBI accession: XM_001792381.1) were generated via homologous recombination (*pnvea-23* and *pnvea-25*) in the *P. nodorum* SN15 background (**Supplementary Fig. S1**). Mutant strains *pnvea-23* and *pnvea-25* were phenotypically compared to SN15 to assess the role of PnVeA in the development of the fungus. Strains lacking *PnVeA* had produced a dark secretion when cultivated on agar medium, which was lacking in SN15, and were unable to produce pycnidia and pycnidiospores (**Fig. 2A and B**).

**Figure 2.**
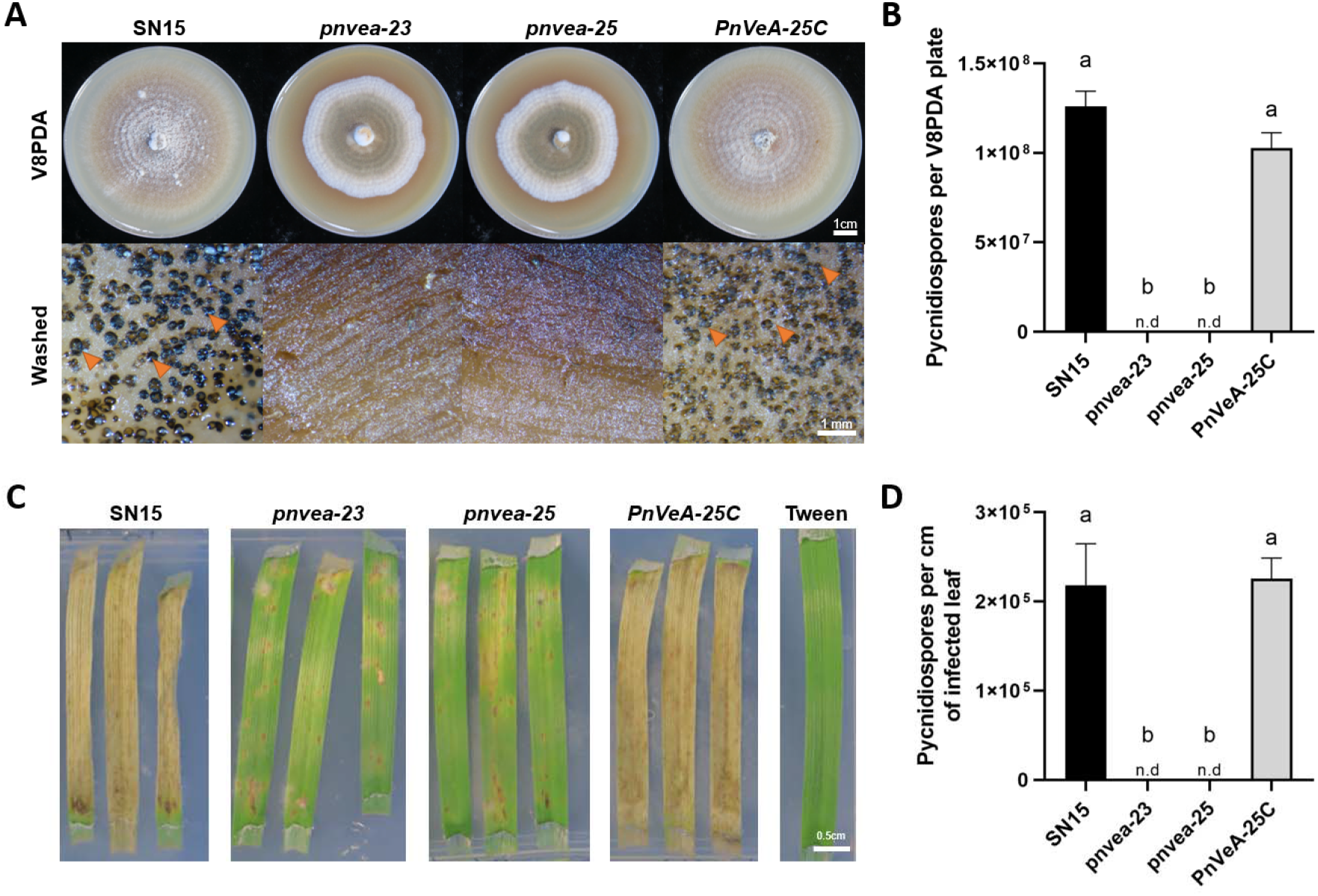
Comparative examination of vegetative morphology and virulence of *P. nodorum pnvea* mutants with SN15. **(A)** Vegetative morphology of *P. nodorum* strains on V8PDA (top row) highlighting differences in pycnidial development (orange arrows) (bottom row). **(B)** Pycnidiospore count of *P. nodorum* strains on V8PDA media after 12 days of growth showing the abolishment of pycnidiospore production in *pnvea-23* and *pnvea-25*. **(C)** Wheat leaf assay infection at day 6 post-inoculation showing reduced virulence of knockout mutants on wheat cultivar Axe. Tween was used as a negative control. **(D)** Pycnidiospore count of *P. nodorum* strains normalized to the length of infected leaf tissue after 10 days, showing the abolishment of spore production in *pnvea-23* and *pnvea-25*. For **(B)** and **(D)**, ANOVA with the Tukey-Kramer test was used to compare the means of pycnidiospore counts (*p* ≤ 0.05, *n* = 3). Different letters above the bars indicate statistical significance between strains. Strains without detectable pycnidiospores are indicated by “n.d.” for “not detected”.

To determine whether *PnVeA* affected virulence *in-planta*, *pnvea-23* and *pnvea-25* were compared to SN15 using a wheat leaf infection assay. Both *pnvea-23* and *pnvea-25* were highly reduced in virulence and unable to produce pycnidia (**Fig. 2C and D; Supplementary Fig. S2**). Microscopic analysis revealed *pnvea-25* was capable of host penetration attempt via the stomata and epidermis comparable to SN15 (**Supplementary Fig. S3**).

To confirm that the phenotype of mutants is the result of the deletion of *PnVeA*, a gene complementation mutant was generated using the *pnvea-25* background (*PnVeA-25C*). Phenotypic analyses indicated that the *P. nodorum PnVeA-25C* is comparable to SN15 (**Fig. 2**).

### RNA Sequencing revealed that *PnVeA* regulates necrotrophic effectors and secondary metabolite genes

As the deletion of PnVeA affects vegetative morphology and virulence, we then used a comparative RNA-Seq approach to investigate the expression profile in SN15 and *pnvea-25* under *in vitro* growth to identify genes responsible for the observed phenotypes. A principal component (PC) analysis of the SN15 and *pnvea-25* RNA-Seq data revealed a clear segregation between SN15 and *pnvea-25* consistent across replicates (**Fig. 3A**). PC1 accounted for 85% of the total variation while PC2 accounted for 12% and this discriminated SN15 from *pnvea-25* (**Fig. 3A**). The *in-vitro* RNA-Seq analysis revealed 5052 differentially expressed (DE) genes between *pnvea-25* and the SN15 with 2898 up-regulated and 2154 down-regulated genes (**Supplementary data S1**). The putative orthologs *PnVelB* and *PnVosA* were significantly down-regulated with a *Log_2_*fold change (LFC) of −1.35 and −5.54, respectively, while *PnVelC* expression remained unaltered.

**Figure 3.**
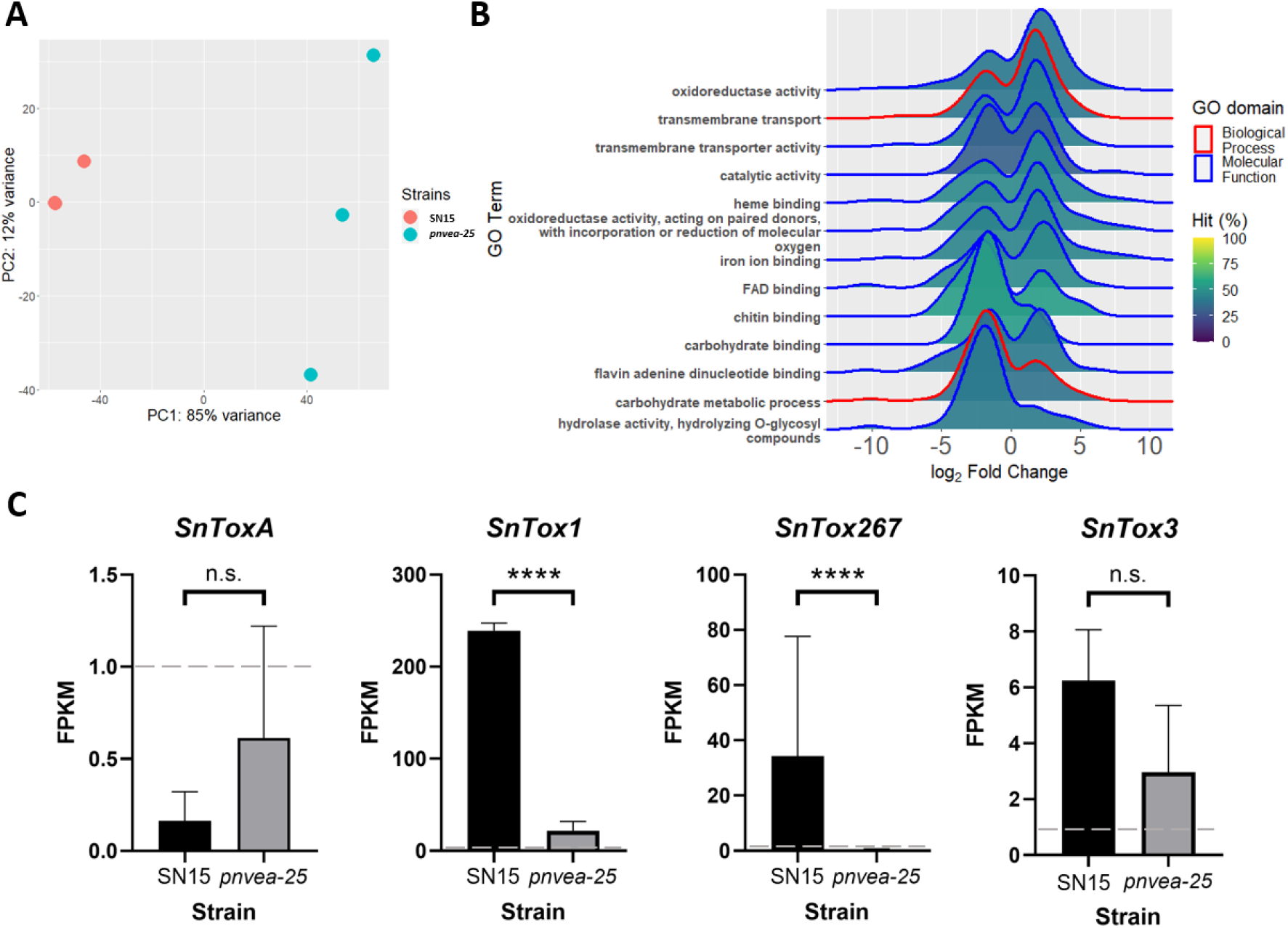
Comparative RNA-Seq analysis between SN15 and *pnvea-25*. **(A)** Principal component analysis (PCA) of transcriptomes of SN15 and *pnvea25*. Three biological replicates were used. Two SN15 biological replicates overlapped on the plot resulting in only two visual data points. **(B)** A ridge plot illustrating the enriched gene ontology (GO) terms between SN15 and *pnvea-25* plotted against the LFC of genes within the GO terms. The shading represents the percentage of genes in the GO term that was enriched. **(C)** The expression profile of necrotrophic effectors *SnToxA*, *SnTox1*, *SnTox267* and *SnTox3* in the RNA-Seq analysis. Horizontal dashed grey lines at one fragment per kilobase of transcript per million mapped reads (FPKM) indicate the lowest threshold at which a gene is considered expressed. Asterisks indicate significant difference while n.s. is not significant according to Wald test with the Benjamini-Hoschberg adjustment, p < 0.0001 (****).

The enrichment of gene ontology (GO) terms within DE genes was analysed to assess the nature of the gene expression changes between SN15 and *pnvea-25*. This revealed that overall, PnVeA down-regulates transport within the cell and up-regulates processes related to carbohydrates (**Fig. 3B**). Major enriched genes up-regulated in *pnvea-25* encoded for oxidoreductase activity (GO:0016491, 138 out of 502 genes), transmembrane transport (GO:0055085, 142 out of 561 genes) and transmembrane transporter activity (GO:0022857, 111 out of 420 genes). Notably, 21 out of 142 down-regulated transmembrane transport genes were predicted to be transporters associated with nitrogen assimilation (**Supplementary data S2**). A similar number of genes in catalytic activity (GO:0003824) and various GO terms related to binding (GO:0020037, GO:0005506, GO:0071949, GO:0008061, GO:0030246, GO:0050660) were equally both up and down-regulated between SN15 and *pnvea-25*. Genes that are associated with carbohydrate metabolic processes (GO:0005975 – 58 out of 229 genes) and hydrolase activity (GO:0004553 – 46 out of 140 genes) were enriched and predominantly down-regulated in *pnvea-25*. Notably, 35 of the 46 down-regulated genes in the GO term hydrolase activity (GO:0005975) were putative plant cell wall depolymerases (**Supplementary data S3**).

*SnToxA*, *SnTox1*, *SnTox267* and *SnTox3* are well-characterised necrotrophic effector-coding genes (Friesen et al. 2006; Friesen et al. 2008; Liu et al. 2004; Richards et al. 2022). In addition, it is hypothesised that *P. nodorum* possesses additional undiscovered effector genes (Jones et al. 2021). These effector genes were then examined in the RNA-Seq analysis between SN15 and *pnvea-25* to explore a connection with known virulence factors. *SnTox1* and *SnTox267* were significantly down-regulated while the expression of *SnToxA* and *SnTox3* was not altered during *in-vitro* growth (**Fig. 3C**). Three effector candidates, SNOG_13888, SNOG_42372 and SNOG_10241 were significantly down-regulated (**Table S1**).

### Deletion of *PnVeA* resulted in the loss of coordinated expression of major necrotrophic effector genes during infection

Deregulation of effector genes in the RNA-Seq prompted an analysis of their expression during host infection. Therefore, digital droplet PCR (ddPCR) was utilised and showed that *pnvea-25* had a significantly different profile of effector gene expression compared to the SN15 and *PnVeA-25C* (**Fig. 4**). Although, the expression *SnToxA* did not significantly change without PnVeA (**Fig. 4A**). In contrast, *SnTox1* expression in *pnvea-25* was higher than in SN15 and *PnVeA-25C* at 10 days post infection (**Fig. 4B**). Interestingly, *pnvea-25* expression of *SnTox3* was increased from six days to 10 days post inoculation *in-planta* compared to SN15 and *PnVeA-25C* (**Fig. 4C**). Furthermore, *SnTox267* expression in *pnvea-25* was significantly higher than in SN15 and *PnVeA-25C in-planta* at 10 days post inoculation (**Fig. 4D**). Overall, *pnvea-25* expressed the effector genes at higher levels than SN15 and *PnVeA-25C* at 10 days post-inoculation, excluding *SnToxA* where expression remained unchanged *in-planta*. This corresponds to pycnidiation in the infection cycle of *P. nodorum*, which was abolished in *pnvea-25* and also where effector expression is typically reduced (IpCho et al. 2010).

**Figure 4.**
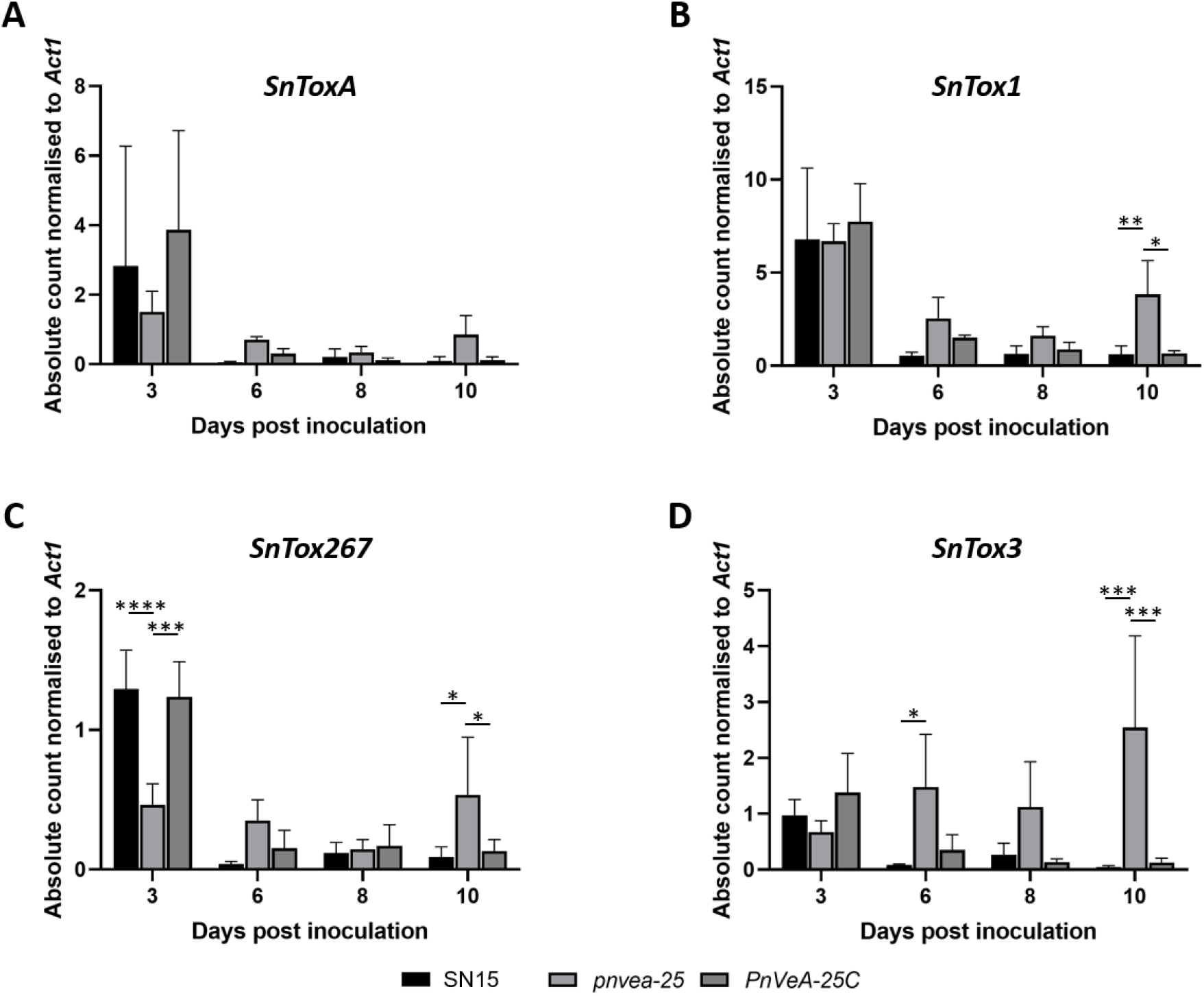
Digital droplet PCR of necrotrophic effector genes revealed that PnVeA coordinates *SnTox1*, *SnTox3* and *SnTox267* expression *in-planta*. The *in-planta* gene expression profiles of effector genes, (**A**) *SnToxA*, (**B**) *SnTox1*, (**C**) *SnTox267* and (**D**) *SnTox3*, for the wildtype strain (SN15), *PnVeA* deleted mutant (*pnvea-25*) and complemented mutant (*PnVeA-25C*) were measured at three-, six-, eight- and 10-days post infection. Asterisks indicate a significant difference in mean within a time point between strains according to a one-way ANOVA with the Tukey-Kramer test post-hoc (p ≤ 0.05, n = 3), p < 0.05 (*), < 0.01 (**), < 0.001 (***) and < 0.0001 (****).

### PnVeA is associated with the regulation of putative secondary metabolite gene clusters

The identification and characterisation of PnVeA presents an opportunity to further investigate SMs in *P. nodorum* at a global regulatory level based on the current understanding of Velvet regulation of SMs in other fungi. In *A. flavus* and the corn pathogen *Fusarium verticillioides*, the VeA orthologs are regulators of secondary metabolism (Duran et al. 2007; Myung et al. 2009) and since SM gene clusters have been previously discovered in *P. nodorum* (Chooi et al. 2014), the expression of these clusters was analysed. Five putative polyketide synthase (PKS) gene clusters, identified using antiSMASH (Chooi et al. 2014), showed differential expression patterns between SN15 and *pnvea-25* in the RNA-seq analysis (**Fig. 5**).

**Figure 5.**
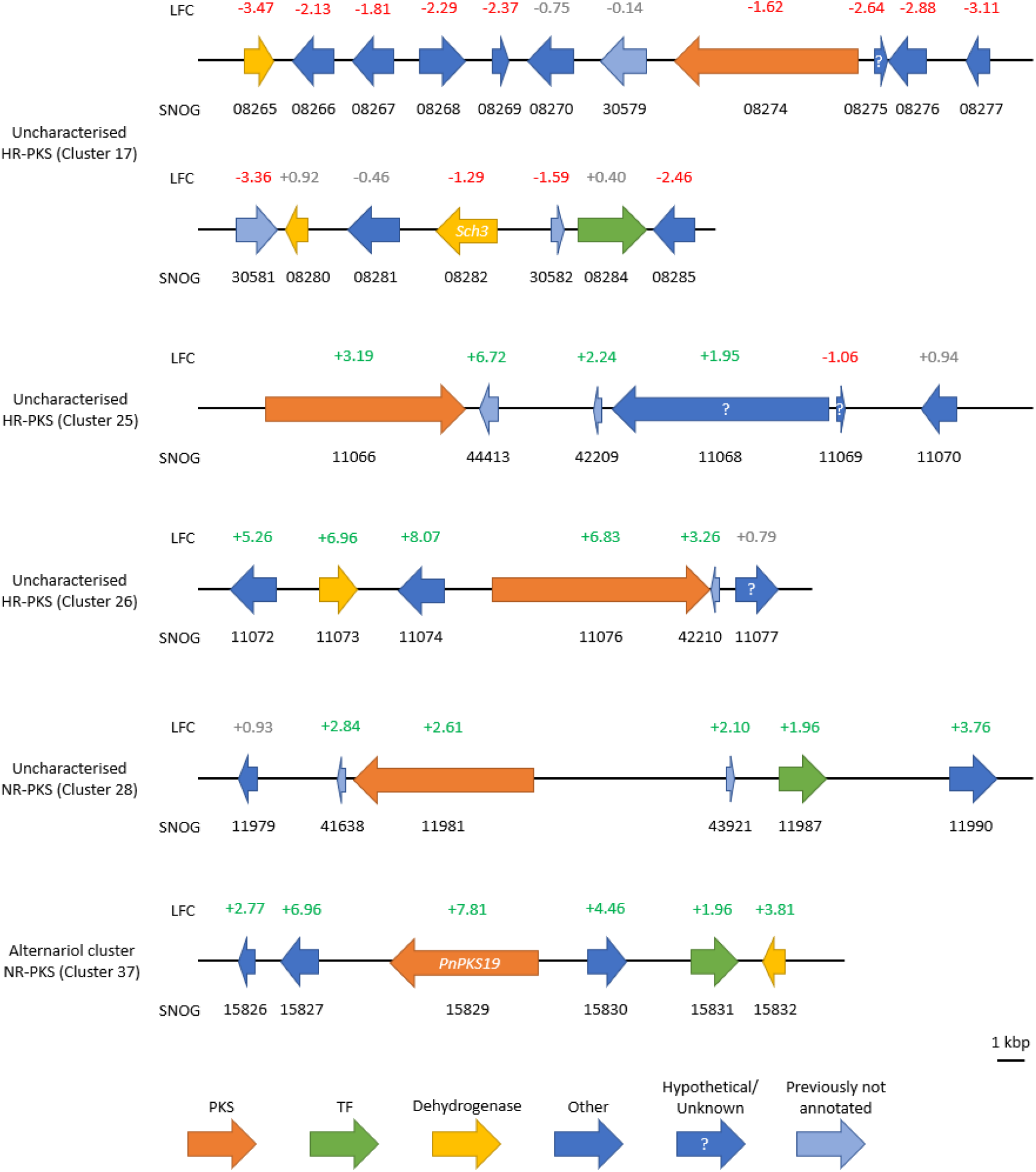
Differential expression of differentially regulated SM gene clusters in *P. nodorum* (Chooi et al. 2014). The cluster designation is by Chooi et al. (2014). *In-vitro* RNA-Seq analysis compared the expression between wildtype SN15 and *PnVeA* deleted strain (*pnvea-25*). *Log_2_* fold change (LFC) is coloured according to its significance: green is significantly up-regulated in *pnvea-25*, red is significantly down-regulated in *pnvea-25* and grey is not significant. SNOG is the gene locus names. “PKS” are polyketide synthase genes, “TF” are transcription factors, “Other” are genes of various miscellaneous functions and “Previously not annotated” are genes within a cluster but was not present in the genome annotation used by Chooi et al. (2014).

Cluster 17 contains the highly reducing PKS (HR-PKS) gene SNOG_08274. Out of the 18 predicted genes in this cluster, 13 were significantly down-regulated, including the backbone gene. antiSMASH comparative gene cluster analysis suggests that Cluster 17 can produce an SM similar to the naphthopyrone YWA1, a conidial pigment intermediate in the opportunistic human pathogenic fungus *A. fumigatus* (Piras et al. 2021). Furthermore, SNOG_08274 has been previously associated with pigment production (Chooi et al. 2014).

Two adjacent SM gene clusters (Cluster 25 and Cluster 26) containing putative HR-PKS genes (SNOG_11066 and SNOG_11076, respectively) were up-regulated in *pnvea-25* mutant. While Cluster 25 has been associated with pyranonigrin E production (Awakawa et al. 2013), antiSMASH analysis has linked Cluster 26 to the macrolide AKML (Morishita et al. 2020). Genes in Cluster 28, which contains a NR-PKS gene SNOG_11981, were also up-regulated. Of the six genes predicted within Cluster 28, five were up-regulated. SNOG_11981 has been linked to melanin production (Fulton et al. 1999).

Notably, of the five DE PKS gene clusters, one (NR-PKS; *PnPKS19*) is involved in the production of alternariol, a previously characterized mycotoxin (Chooi et al. 2015). To assess the correlation between RNA-seq results and SM production, particularly the abundance of alternariol, the SM profiles of all strains were analysed through LC-DAD-MS. *PnVeA* deleted strains (*pnvea-23* and *pnvea-25*) displayed a new peak **1** (m/z 257 [M-H]-) (Chooi et al. 2015), also present in lower quantities in *PnVeA-25C*, but undetectable in the control strain SN15 (**Fig. 6A**). Besides having a similar mass, the compound UV-vis spectrum and molecular formula were consistent with alternariol (**Fig. 6B and C, Supplementary Fig. S4**).

**Figure 6:**
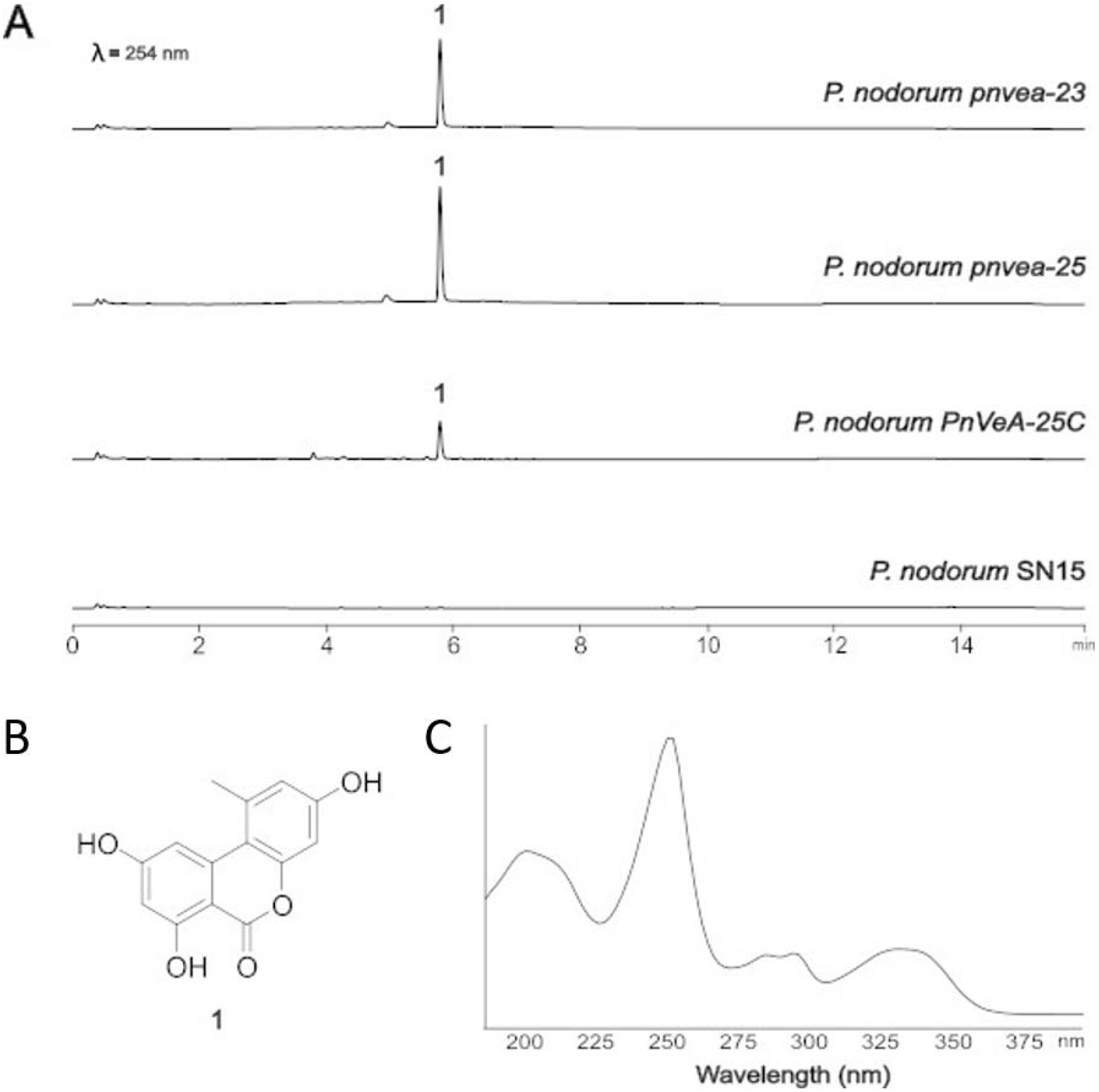
The deletion of *PnVeA* resulted in a modified metabolic profile. **(A)** LC-DAD (DAD 254 nm) chromatograms showing the metabolomic profiles of *P. nodorum* SN15, *PnVeA*, *pnvea* and *PnVeA-25C* strains grown on V8PDA. Deletion of *PnVeA* led to increased production of **1** (alternariol), the structure of which is depicted in **B**. **(C)** Alternariol UV-vis spectrum.

Additionally, two uncharacterized compounds, **2** (m/z 378 [M+H]+) and **3** (m/z 392 [M+H]+), without apparent UV absorption were detected in SN15 and *PnVeA-25C*, but were absent in *pnvea-23* and *pnvea-25* (**Supplementary Fig. S5**).

## Discussion

The Velvet TFs are known to function as global regulators involved in diverse functions in filamentous fungi (Calvo et al. 2016; John et al. 2021). Until now, their regulatory role had not been investigated in the NE-producing pathogen *P. nodorum*. Using a gene deletion approach, we highlight several key processes controlled by the VeA ortholog. Phenotypic analysis and RNA sequencing data suggest that PnVeA is required for asexual sporulation, normal effector regulation and SM production.

Additional growth abnormalities in *P. nodorum* were observed in the *PnVeA* deletion mutants such as a high pigmentation and reminiscence of the ‘velvet’-like texture originally described in *A. nidulans* (Käfer 1965). VeA orthologs in other characterised fungi are not required for sporulation but have significantly different asexual and sexual spore counts than the wild-type strain (Calvo et al. 2016). PnVeA was uniquely required for sporulation as strains without a functional *PnVeA* gene had complete abolishment of pycnidia and pycnidiospore production. This unusual phenotype was observed in both *in-vitro* condition and during infection indicating that PnVeA has a more pivotal role in asexual sporulation in *P. nodorum* than VeA orthologs in other characterised fungal species such as the fungal pathogens: *A. flavus*, *B. cinerea* and *M. oryzae* (Eom et al. 2018; Kim et al. 2014; Müller et al. 2018; Schumacher et al. 2012). Due to the difficulty of obtaining *P. nodorum* ascospores under laboratory conditions (Solomon et al. 2006), the impact of PnVeA on sexual reproduction could not be assessed.

Comparative RNA-Seq analysis revealed that PnVeA has a global role in gene expression, including dysregulation of *P. nodorum* carbohydrate/chitin-associated genes, iron/FAD metabolism, secondary metabolite biosynthesis as well as NE gene expression. Leaf infection assays using *pnvea-23* or *pnvea-25* demonstrate reduced virulence despite retaining the capacity to penetrate the host tissue. The effector gene expressions were investigated further during infection with ddPCR, which showed variable profiles as a result of PnVeA presence or absence. It is likely that down-regulation in extracellular hydrolases, perturbation in biochemical processes or down-regulation of other undiscovered effectors can be attributed to the reduction in virulence on wheat that cannot be overcome with NEs. With down-regulation of putative plant cell wall degrading enzyme (CWDE) genes observed in *pnvea-25*, the mutant may have perturbed ability to breakdown cell walls and therefore unable to colonise the host tissue effectively. *pnvea-25* may in turn compensate for the reduced ability to hydrolyse cell walls by increasing the expression of membrane transporter genes. Reduced expression of hydrolases and increased expression of transporters were seen in the RNA-Seq analysis of a *P. nodorum* mutant lacking PnPf2, a TF involved in virulence and required for the expression of *SnToxA* and *SnTox3* (Jones et al. 2019; Rybak et al. 2017). As the expression of *PnPf2* was down-regulated by 1.7-fold in *pnvea-25* compared to SN15, there may be an *in-vitro* connection. A previous study investigating the essential TF *PnCon7* reveals a reduction in virulence during infection in *PnCon7*-silenced mutant (Lin et al. 2018). *PnCon7* is a fundamental developmental regulator but also directly controls *SnTox3* expression, as well as influencing *SnTox1* expression (Lin et al. 2018). *PnCon7* was significantly down-regulated in *pnvea-25* compared to SN15 indicating that there may be a connected role *in-vitro* as a similar decrease in expression of *SnTox1* and *SnTox3* was observed in *pnvea-25* although, *SnTox3* expression was not statistically significant. Furthermore, *SnToxA* was not expressed in either SN15 nor *pnvea-25 in-vitro* and this was consistent with previous literature (Rybak et al. 2017; Tan et al. 2015).

Interestingly, *pnvea-23* or *pnvea-25* showed no chlorosis nor necrosis during early infection despite the expression of all the effector genes of *pnvea-25 in-planta* being comparable to SN15 and *PnVeA-25C*, except for *Tox267* at three days post inoculation. The reduced chlorosis and necrosis of *pnvea-23* and *pnvea-25* may be attributed to the reduced expression of CWDE genes. The infection by *pnvea-23* and *pnvea-25* showed necrotic symptoms at 10 days post inoculation and the expression of effector genes was much higher than SN15 or *PnVeA-25C*. This suggests that PnVeA may be involved in the coordination of additional factors necessary for NE gene regulation rather than directly regulating the expression of NE genes themselves. There may be an undiscovered role of PnVeA related to detection of wheat environmental cues to enable infection, reflected by the delayed expression of NE genes and subsequent appearance necrotic symptoms.

In addition to *SnTox1* and *SnTox267*, the expression of 13 NE candidates identified by Jones et al. (2021) was also altered *in-vitro*. Of these, the three effector candidates with the highest negative fold change in *pnvea-25* were SNOG_10241, SNOG_13888 and SNOG_42372. SNOG_10241 is a homolog to Zt6, an effector found in the wheat pathogen *Zymoseptoria tritici* that is phytotoxic and possesses antimicrobial properties (Kettles et al. 2018). Interestingly, the deletion of *Zt6* did not affect the virulence of *Z. tritici* on wheat (Kettles et al. 2018). The characterised effector closest to SNOG_13888, MoBas4, is associated with the suppression of host defence by *M. oryzae* (Sharpee et al. 2017). SNOG_42372 homologous to chitin-binding effectors of the LysM family which conceal pathogens by outcompeting chitin-binding proteins from the host that trigger an immune response (Kombrink and Thomma 2013). Silencing and deletion of LysM effector genes result in reduced virulence such as the effector LtLysM1 in the wide host range pathogen *Lasiodiplodia theobromae* (Harishchandra et al. 2020) and Vd2LysM in *Verticillium dahliae*, a tomato pathogen (Kombrink et al. 2017). The characterised orthologs of SNOG_10241, SNOG_13888 and SNOG_42372 are either phytotoxic or used by the pathogens to evade host immune response and therefore these putative orthologs may facilitate the phytopathogenicity of *P. nodorum*.

Alternariol is a mycotoxin found in *P. nodorum* and in the members of the fungal genus *Alternaria*, including the broad host phytopathogen *Alternaria alternata* (Ostry 2008; Tan et al. 2009). Using LC-DAD-MS, PnVeA appeared to negatively regulate the production of alternariol. This is in direct contrast to what has been observed in *A. alternata,* where VeA was a positive regulator of alternariol production (Wang et al. 2022). The previously characterised PKS gene, *PnPKS19* is required to produce alternariol in *P. nodorum* (Chooi et al. 2015). The orthologs of the *PnPKS19-*containing cluster also positively regulate alternariol production in *A. alternata* (Chooi et al. 2014; Wenderoth et al. 2019). Deletion of VeA in *A. alternata* resulted in reduced expression of the *PnPKS19* ortholog (*pksI*) (Wang et al. 2022) while the RNA-Seq DE gene analysis between SN15 and *pnvea-25* indicated that *PnPKS19* was the ninth most up-regulated gene across the whole genome. It is unknown why the VeA-orthologs regulate alternariol production differently between *A. alternata* and *P. nodorum*. Short-chain dehydrogenase 1 (Sch1) is a negative regulator of alternariol production (Tan et al. 2009). *Sch1* deletion results in the accumulation of alternariol likely through pleiotropic effects (Tan et al. 2009). Analysis of *Sch1* expression indicates that it is unchanged in *pnvea-25*. This suggests that the synthesis of alternariol is directly attributed by PnPKS19 overcomes the suppression of alternariol production by Sch1. Three interesting uncharacterised PKS genes include SNOG_08274 (Cluster 17), SNOG_11076 (Cluster 26) and SNOG_11981 (Cluster 28). The identity of the metabolite product associated with the SNOG_08274 gene cluster is currently unknown. However, the short-chain dehydrogenase gene *Sch3* (SNOG_08282) is located within the cluster (Casey et al. 2010) (Fig. 5). Functional characterisation indicates that *Sch3* is required for asexual sporulation. Chooi et al. (2014) proposed that Cluster 17 shared the closest BLAST hit to the PKSN gene cluster associated with pigmentation of sexual fruiting bodies of the phytopathogenic fungus, *Nectria haematococcoa* (Casey et al. 2010; Chooi et al. 2014; Graziani et al. 2004). As the most similar antiSMASH hit for SNOG_08274 indicated a PKS with a conidial pigment intermediate (Piras et al. 2021), there is a possible connection between the non-sporulating phenotype of *pnvea-23* and *pnvea-25* and the down-regulation of the SNOG_08274-containing gene cluster.

SNOG_11981 was predicted to be involved in the production of melanin (Fulton et al. 1999). An increase in the expression of the SNOG_11981-containing gene cluster may be attributed to the hyperpigmentation phenotype of *pnvea-23* and *pnvea-25*. A fourth cluster, with the HR-PKS gene SNOG_11066, was also DE in *pnvea-25*. However, it was previously shown that the SNOG_11066 gene itself is not expressed in *P. nodorum* (Chooi et al. 2014; Ipcho et al. 2012). The pyranonigrin E gene cluster in *A. niger*, containing the most similar PKS to SNOG_11066, was also found to be not expressed (Awakawa et al. 2013).

Interestingly, in the LC-DAD-MS analysis of SM extracts from SN15 and *PnVeA-25C*, two compounds without UV absorption were detected. These compounds were not detected in *pnvea-23* and *pnvea-*25. The cluster containing SNOG_08274 was the sole one that appeared down-regulated. Whether these molecules are associated with the SNOG_08274 cluster xx requires further experimental validation. Although tools like antiSMASH can predict SM-related genes for various classes of molecules, the biosynthetic origin of many compounds remains to be determined (Yee et al. 2023). Consequently, molecules we detected might be products of genes that lie outside the gene clusters recognised by antiSMASH or result from unclustered biosynthesis pathways. Furthermore, our findings suggest that VeA is a crucial global regulator, suppressing alternariol production while potentially influencing the synthesis of other molecules.

Over 3000 genes are potentially directly regulated by VeA in *A. nidulans* (Moon et al. 2023), so it is likely similar for PnVeA. Therefore, further characterisation of PnVeA is required to determine its direct targets.

This study has characterised PnVeA, the ortholog of a fungal-specific global regulator VeA. It was discovered that PnVeA appears to regulate growth, pigmentation and sporulation, as well as both effector expression and SM production. Uniquely, in *P. nodorum*, PnVeA is essential for sporulation. Furthermore, PnVeA is a negative regulator of alternariol production and is the first TF in *P. nodorum* observed to putatively regulate the NE gene *SnTox267*. Four uncharacterised SM gene clusters identified to be regulated by PnVeA were observed in this study. Efforts are underway to investigate SMs attributed to these canonical SM gene clusters. In addition, chromatin immunoprecipitation (Park 2009) is being used to uncover direct gene targets of PnVeA.

## Materials and Methods

### Strains and cultures

*P. nodorum* strains used in this study are listed in **Table 1**. All fungal strains were maintained on V8PDA (150 mL/L Campbell’s V8 Juice, 3 g/L CaCO_3_, 10 g/L Difco PDA and 10 g/L Agar) (Solomon et al. 2004). Fungal strains were grown in a 12: 12 hr light: dark cycle at 22°C for 7 to 12 days.

**Table 1:** Fungal strains used in this study.

| Strain | Description | Source |
| --- | --- | --- |
| SN15 | *Parastagonospora nodorum* wildtype | Department of Primary Industries and Regional Development, Western Australia |
| *pnvea-23* | SN15 carrying an in-frame deletion of *PnVeA* | This study |
| *pnvea-25* | SN15 carrying an in-frame deletion of *PnVeA* | This study |
| *PnVeA-25C* | *pnvea-25* complemented with *PnVeA* | This study |

### Identification of Velvet orthologs in *P. nodorum* and phylogenetic tree construction

Velvet proteins VeA (NCBI accession: XP_658656.1), VelB (XP_657967.1), VelC (XP_659663.1) and VosA (XP_659563.1) from *A. nidulans* were used to search for the ortholog in *P. nodorum* reference strain SN15 (Bertazzoni et al. 2021) using the Basic Local Alignment Search Tool blastp (Altschul et al. 1997). The probable orthologs identified were analysed using InterPro (Paysan-Lafosse et al. 2022) to confirm the presence of the Velvet domains. Characterised velvet proteins from *A. nidulans* and three plant pathogenic fungi (*A. flavus*, *B. cinerea* and *M. oryzae*) were retrieved from UniProt (Consortium 2022) and selected to compare against the identified orthologs in *P. nodorum* (Eom et al. 2018; Kim et al. 2014; Müller et al. 2018; Schumacher et al. 2012). An alignment of full length proteins was performed on MEGA11 using the ClustalW algorithm with default settings (Tamura et al. 2021). A neighbour-joining phylogenetic tree was constructed using default settings except with a bootstrap analysis set at 3000 repetitions (Tamura et al. 2021).

### Generation of *PnVeA P. nodorum* mutants

The primers used in this study are listed in **Table S2**. The gene deletion construct of *PnVeA* including a phleomycin selectable marker was assembled using Golden Gate cloning system (Engler et al. 2008) using a modified pUC19 (New England Biolabs, Ipswich, Massachusetts, USA) as the backbone vector (John et al. 2022). The *PnVeA* complementation construct that included a 1,021 bp promoter region, the full intact gene and 290 bp of the terminator sequence was fused to a hygromycin selectable marker via fusion PCR (Hilgarth and Lanigan 2020).

The *P. nodorum* wild-type strain, SN15, was transformed via polyethylene glycol-mediated transformation as previously described (Solomon et al. 2004). *PnVeA*-deleted *P. nodorum* mutant strain, *pnvea-25*, was similarly transformed but the hyphae were used for the inoculation of the starter culture instead of spores due to its non-sporulating phenotype.

Deletion of the gene and subsequent complementation were confirmed using PCR (**Supplementary Fig. S1A and B**). Single copy integration of the constructs was confirmed using a previously developed quantitative PCR method (**Supplementary Fig. S1C**) (Solomon et al. 2008).

### Phenotypic assay

Agar plugs were cut out from the growing edge of each strain with the back of a 200 µL pipette tip and inoculated onto another V8PDA plate and left to grow under the conditions outlined before. Spores were then counted for each strain after 12 days.

### Infection assay

Hyphae from each strain were homogenised in 0.02% Tween solution using metal beads and a ball mill shaken at 30 Hz for 10 s using a Mixer Mill MM400 (Retsch, Haan, Germany). The hyphal mixture was diluted to 20 mg/mL by wet weight in 0.02% Tween 20. This hyphal mixture was painted using a paintbrush on 10-day-old wheat seedlings (cv. Axe) embedded in 0.5% benzimidazole water agar and grown as a detached leaf assay (Solomon et al. 2004). After 10 days, the infected leaves were put into 500 μL of water and shaken at 30 Hz for 10 s to dislodge the pycnidiospores. The spores were then counted and normalised to spores per cm of infected leaf tissue.

The staining of infected leaves with Trypan Blue was as previously described (Solomon et al. 2004).

### RNA extraction, sequencing and quality control

SN15 and *pnvea-25* were grown on V8PDA for 12 days. RNA extraction of was performed using an extraction involving TRIzol^TM^ reagent (Invitrogen, La Jolla, USA) and DNase treatment using DNase I (New England Biolabs, Ipswich, USA). Library preparation and sequencing were performed by the Australian Genome Research Facility (https://www.agrf.org.au/). Following the library preparation, 150 bp paired-end stranded sequences were generated on a NovaSeq S4 (Illumina, San Diego, USA). The experiment was performed with three biological replicates. The quality of the raw RNA-Seq reads was assessed in FastQC v0.11.9 (https://www.bioinformatics.babraham.ac.uk/projects/fastqc/). The sequencing adapters were trimmed using Trimmomatic v0.39 (Bolger et al. 2014) and the trimmed paired reads were mapped to the *P. nodorum* SN15 genome (Bertazzoni et al. 2021) with known splice sites using STAR v2.7.3 using default settings (Dobin et al. 2013).

### Determination and functional enrichment of differentially expressed genes in RNA-Seq

The counts of the reads overlapping the annotated gene features in the genome were determined using the featureCount v2.0.0 package in SubRead (Liao et al. 2014). Differentially expressed (DE) genes were determined using the R package DESeq2 v1.36.0 (Love et al. 2014). The criterion for a DE gene was the LFC against the null hypothesis −1 ≤ *LFC* ≤ 1 (*H_a_* = |*LFC*| > 1) with the Benjamini-Hoschberg adjusted *P*-value significance threshold of 0.05.

Gene ontology (GO) terms in DE genes were analysed in Goseq v1.26.0 to determine its overrepresentation (Young et al. 2010). GO terms were assigned using InterProScan where possible (Quevillon et al. 2005). Predicted function of genes in the top five most dysregulated GO terms were added using predictions from UniProt (Consortium 2022). SM cluster genes described by Chooi et al. (2014) that were DE were analysed further on antiSmash 7 fungal version (Blin et al. 2023) with default settings using the SN15 genome (Bertazzoni et al. 2021) as the query.

### Gene expression analysis of effector genes in *in-planta* condition

RNA extraction was similarly performed on *in-planta* samples of all strains in this study. RNA from infected wheat leaves (cv. Axe) three-, six-, eight- and 10-days post inoculation was extracted. Reverse transcription PCR was performed on the sample to generate cDNA using iScript^TM^ cDNA Synthesis Kit (BioRad, Hercules, USA) according to manufacturer’s protocol. A QXDx AutoDG ddPCR system (BioRad) was used to perform ddPCR on biological triplicates of the samples. The expression of *SnToxA*, *SnTox1*, *SnTox267* and *SnTox3* was normalised to the expression of *Act1* in the form of absolute quantification (Lin et al. 2018).

### Secondary metabolite assay

Three plates of each *P. nodorum* strains were grown on V8 PDA were cut into approximately 1 cm x 1 cm pieces and immersed in 25 mL of methanol. This immersion was kept on ice and sonicated using a UW3100 sonicator with a VS70 tip (Bandelin, Berlin, Germany) for 2 min at 10 s: 10 s on: off bursts at 50% amplitude to disrupt the cells. This was then centrifuged at 3900 g for 5 min, the supernatant filtered through a 0.2 μm filter, and then was methanol evaporated off in a vacuum to retain the solid organic extracts. The crude extracts were re-dissolved in methanol for LC-DAD-MS analysis. Chromatographic separation was achieved with a linear gradient of 5–95% acetonitrile-H2O (0.1% (v/v) formic acid) over 10 min, followed by 95% acetonitrile for 3 min, with a flow rate of 0.6 mL/min. The MS data were collected in the m/z range of 100–1000. The experiments were performed using an Agilent 1260 liquid chromatography (LC) system, coupled to a diode array detector (DAD) and an Agilent 6130 Quadrupole mass spectrometer (MS) with an electrospray ionisation source. Chromatographic separation was carried out at 40 °C, employing a Kinetex C18 column (2.6 µm, 2.1 x 100 mm; Phenomenex).

## Acknowledgments

This study was supported by the Centre for Crop and Disease Management, a joint initiative of Curtin University and the Grains Research and Development Corporation under the research grant CUR00023 Project F3.4 awarded to KCT. SM was supported by an Australian Government Research Training Program Scholarship. We would like to acknowledge Dr Huyen T. T. Phan for providing the *SnTox267* qPCR primers.

## Supplementary Information

**Supplementary Figure S1.**
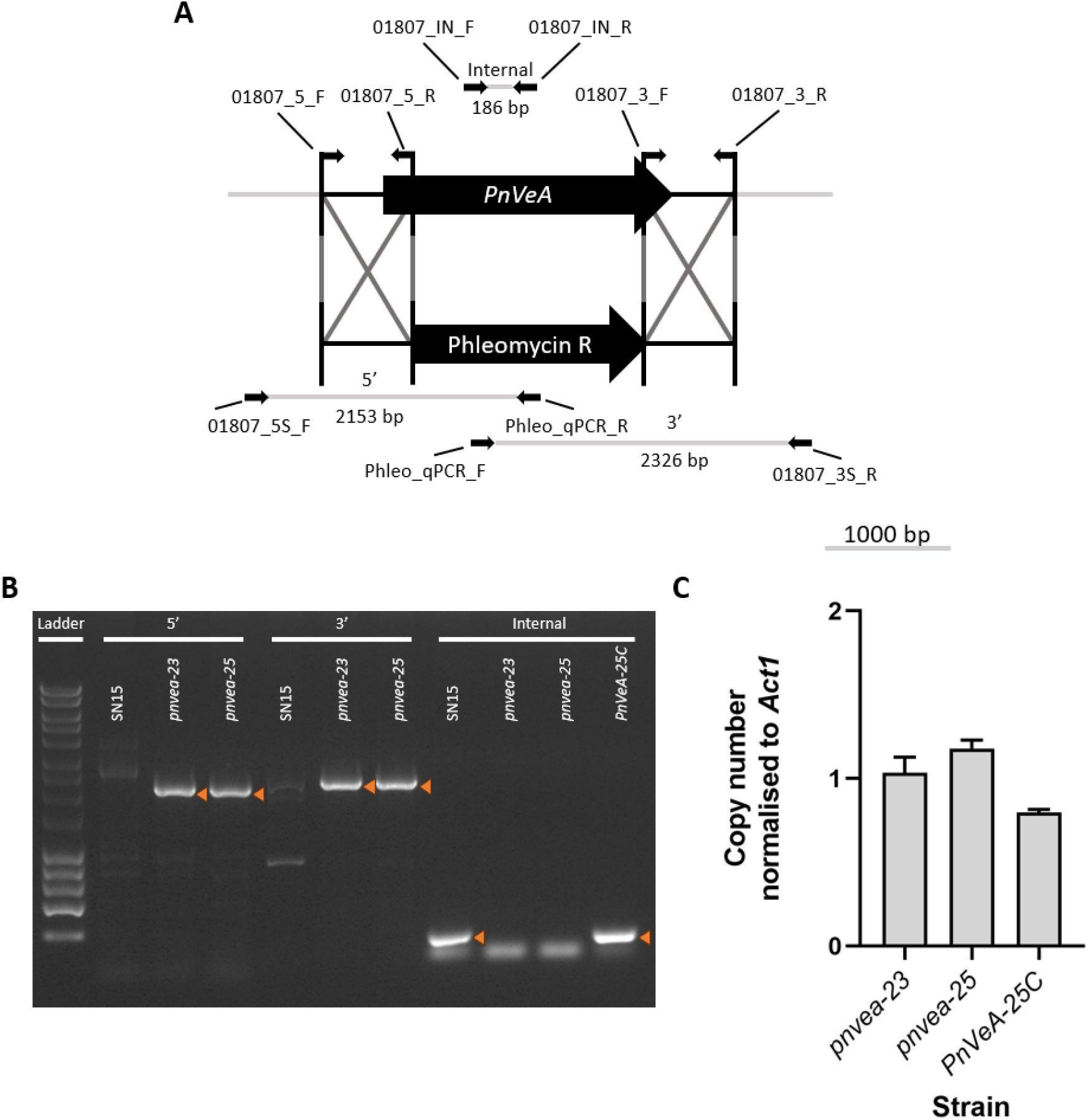
Confirmation of *PnVeA* mutants in this study through PCR and quantitative PCR. **(A)** The construction of *PnVeA* deletion vector and locations of screening primers. **(B)** PCR amplification of screening primers. Amplification of 5’ and 3’ indicate correct integration of the vector. Amplification of “Internal” indicates presence of *PnVeA*. Correct amplifications are indicated by the orange arrow. “Ladder” is the Bioline HyperLadder™ 1 kb marker. **(C)** Copy number of the vectors normalised to a single copy of *Act1* (*n* = 3) (Solomon et al. 2006).

**Supplementary Figure S2.**
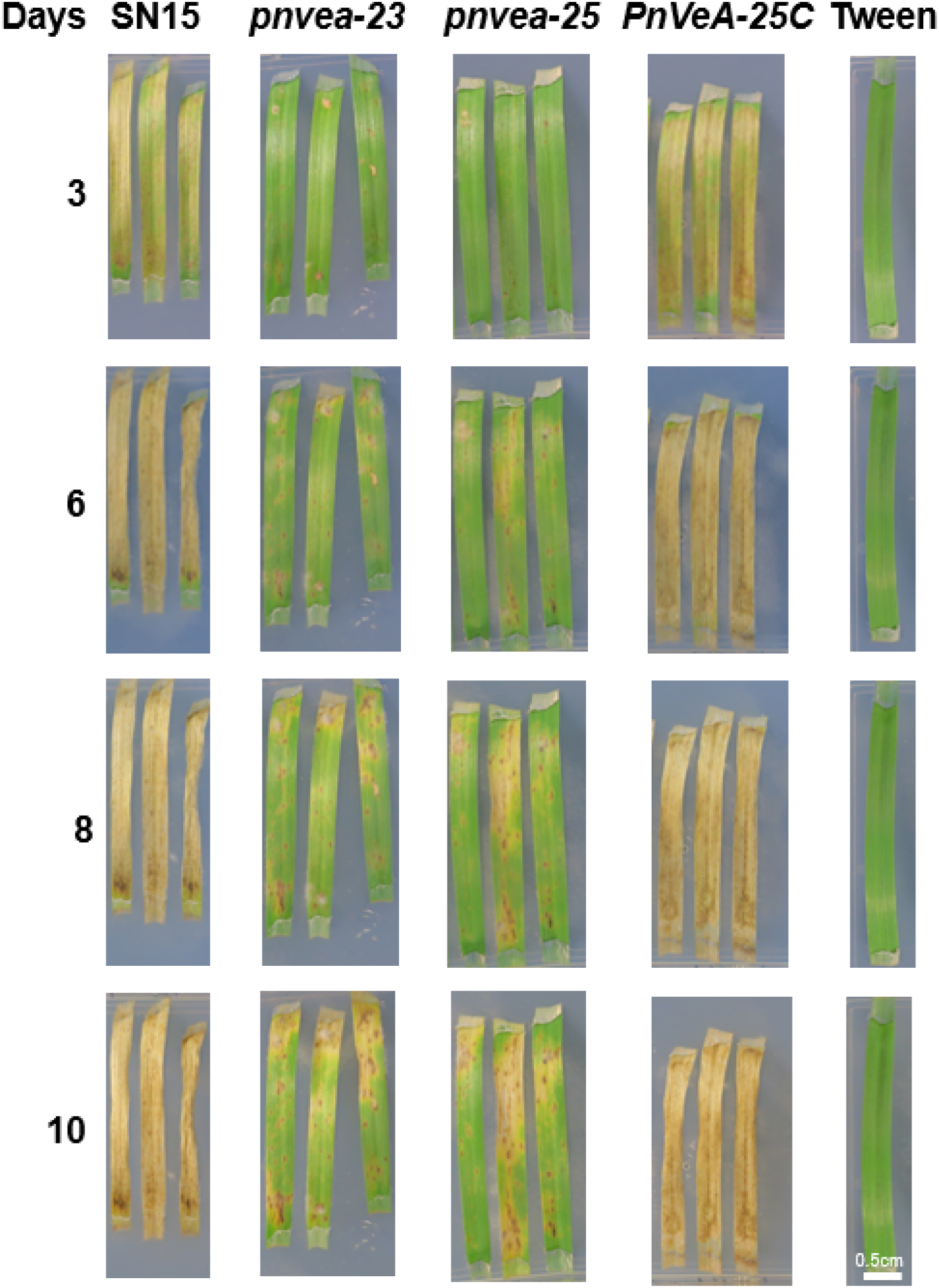
Infection of *Parastagonospora nodorum* strains on wheat (cv. Axe) three-, six-, eight- and 10-days after inoculation. *PnVeA* deleted mutants, *pnvea-23* and *pnvea-25*, showed reduced virulence at every timepoint compared to wildtype (SN15) and *PnVeA* complemented mutant (*PnVeA-25C*).

**Supplementary Figure S3.**
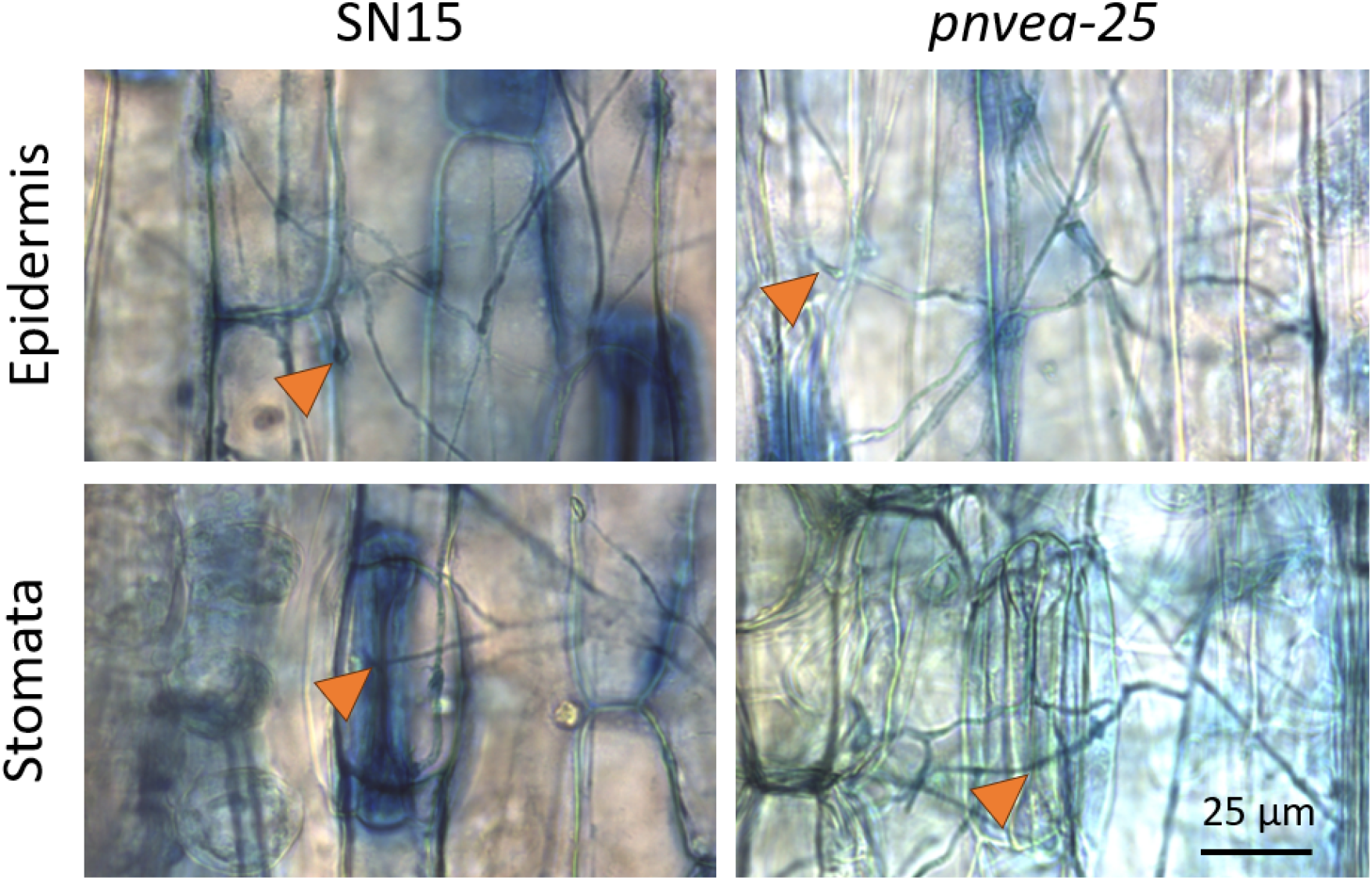
Trypan blue staining of *P. nodorum* strains infecting wheat (cv. Axe) 10 days after inoculation shows no difference in hyphal morphology between the wildtype (SN15) and a mutant with a deletion of *PnVeA* (*pnvea-25*). Orange arrows indicate the location of penetration in either the epidermis (first row) or stomata (second row) of the wheat leaves.

**Supplementary Figure S4.**
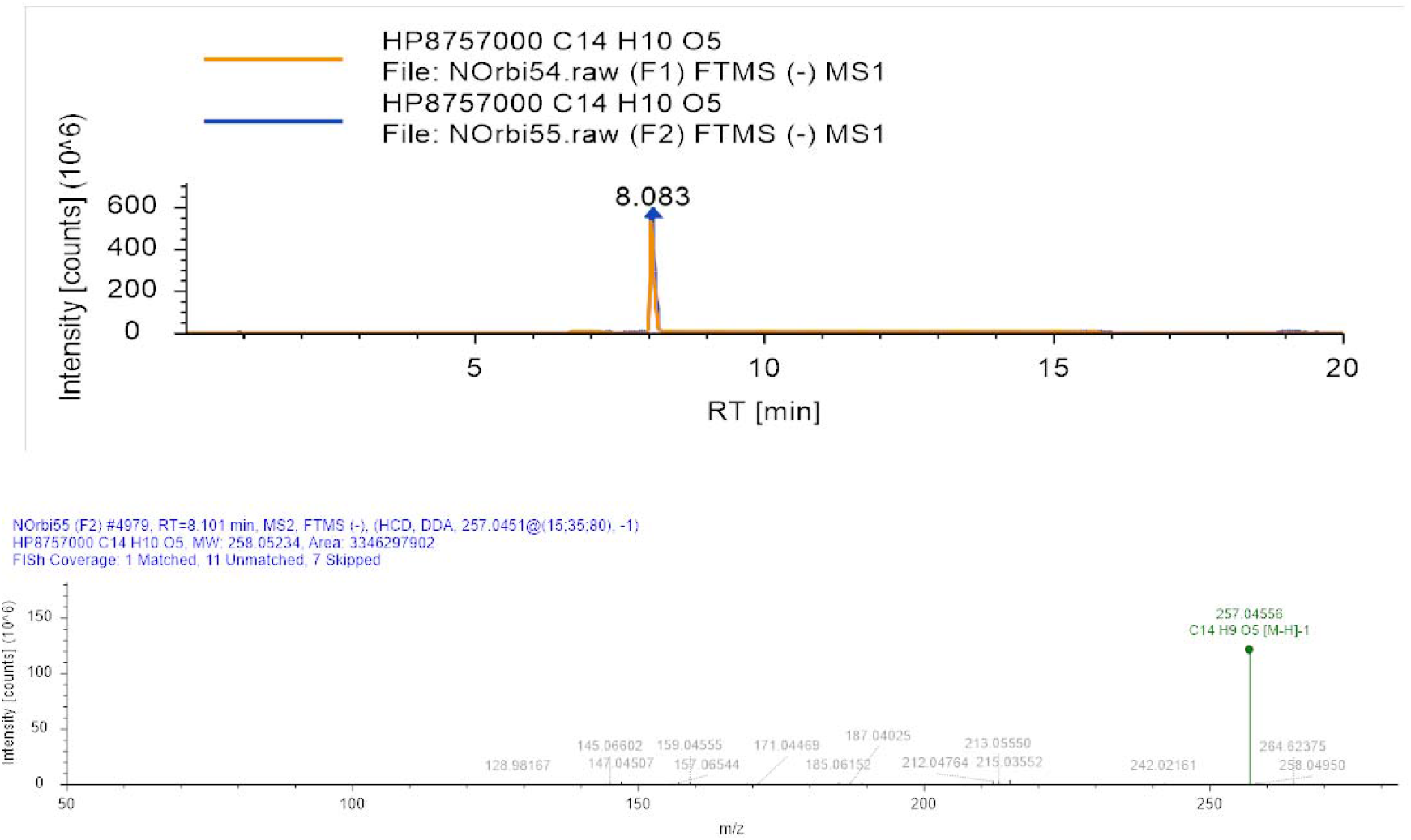
High resolution-electrospray ionization mass spectrometry analysis of compound 1 confirming the atomic formula of alternariol (C14H9O5).

**Supplementary Figure S5.**
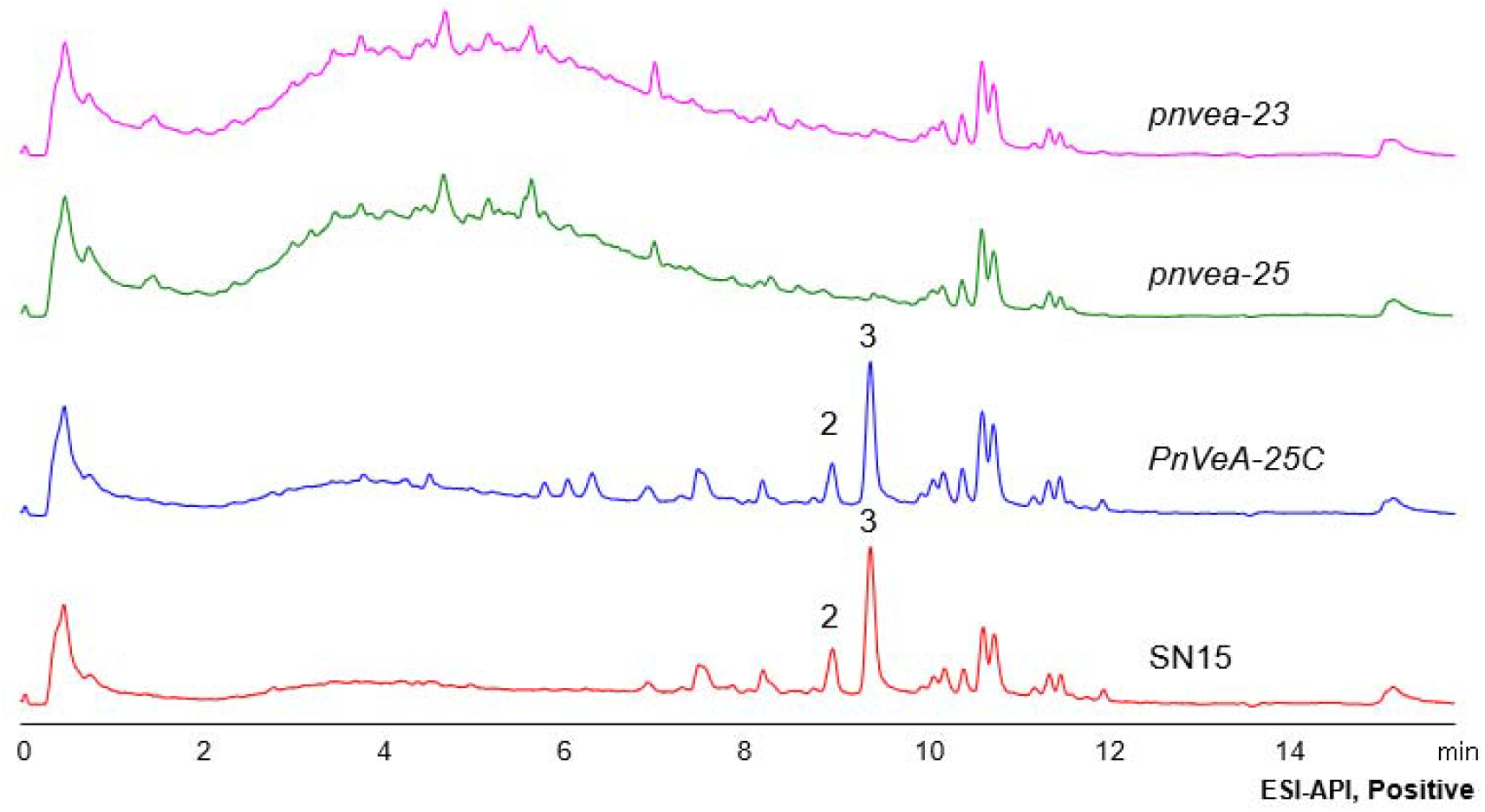
A LC-DAD-MS chromatogram showing the metabolomic profiles of *P. nodorum* wildtype SN15 strain and *PnVeA* deleted mutants (*pnvea-23* and *pnvea-25*) grown on V8PDA. Deletion of the *PnVeA* in *pnvea-23* and *pnvea-25* resulted in decreased production of compounds 2 and 3 (observed in the SN15 and *PnVeA-25C*). A broad peak peaking at four to five minutes was detected in *pnvea-23* and *pnvea-25* but absent in both SN15 and *PnVeA-25C*.

**Table S1.**
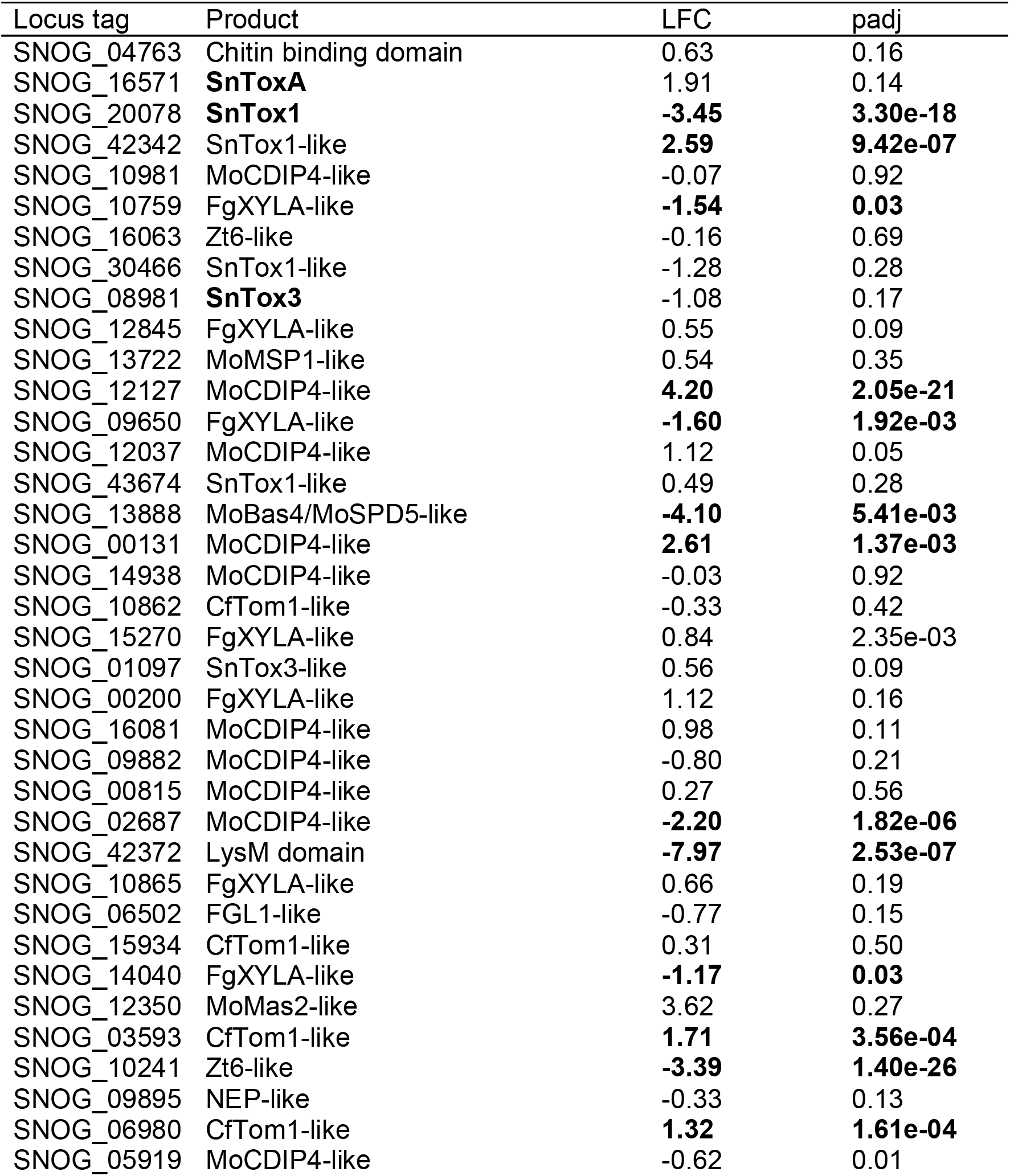
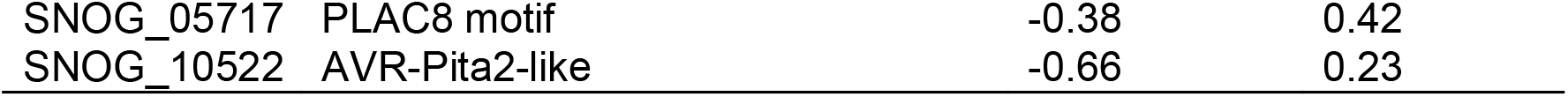
Differential expression of effector candidate genes found in the genome of *Parastagonospora nodorum* SN15 isolate (Jones et al. 2021). RNA-Seq analysis compared the expression between SN15 and *PnVeA* deleted mutant (*pnvea-25*) *in-vitro*. Bolded product name indicates that it has been characterised while bolded *Log*_2_ fold change (LFC) and adjusted p-value (padj) indicate significant differential expression between SN15 and *pnvea-25*.

**Table S2.**
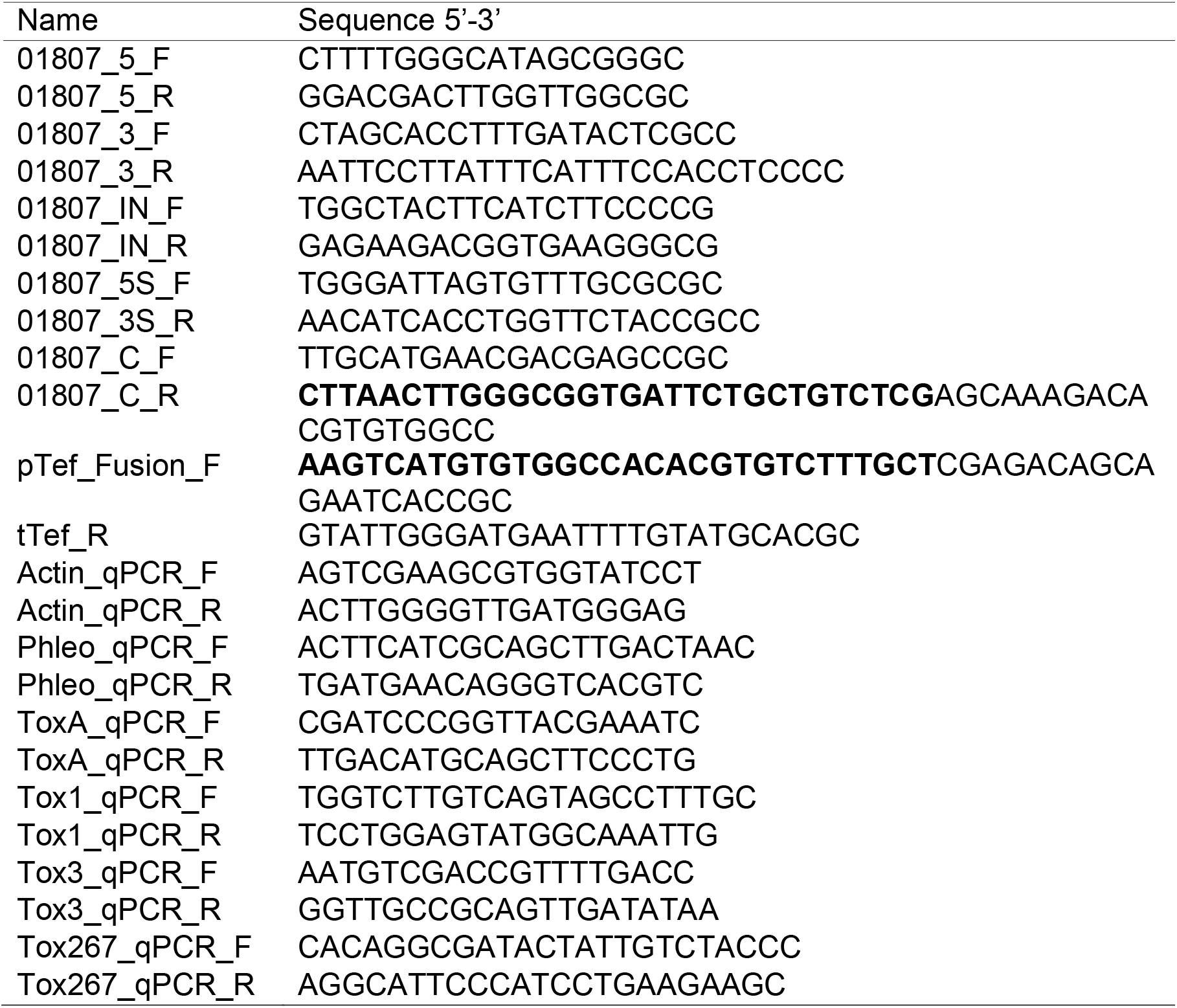
Primers used in this study. Bolded sequences represent overlaps used for fusion PCR.

## References

Ahmed, Y. L., Gerke, J., Park, H.-S., Bayram, Ö., Neumann, P., Ni, M., Dickmanns, A. et al. 2014. The Velvet Family of Fungal Regulators Contains a DNA-Binding Domain Structurally Similar to NF-κB. PLOS Biology 11:e1001750.

Altschul, S. F., Madden, T. L., Schäffer, A. A., Zhang, J., Zhang, Z., Miller, W., and Lipman, D. J. 1997. Gapped BLAST and PSI-BLAST: a new generation of protein database search programs. Nucleic Acids Research 25:3389–3402.

Awakawa, T., Yang, X.-L., Wakimoto, T., and Abe, I. 2013. Pyranonigrin E: A PKS-NRPS hybrid metabolite from Aspergillus niger identified by genome mining. ChemBioChem 14:2095–2099.

Bayram, Ö., and Braus, G. H. 2012. Coordination of secondary metabolism and development in fungi: the velvet family of regulatory proteins. FEMS Microbiology Reviews 36:1–24.

Bertazzoni, S., Jones, D. A. B., Phan, H. T., Tan, K. C., and Hane, J. K. 2021. Chromosome-level genome assembly and manually-curated proteome of model necrotroph Parastagonospora nodorum Sn15 reveals a genome-wide trove of candidate effector homologs, and redundancy of virulence-related functions within an accessory chromosome. BMC Genomics 22:382.

Blin, K., Shaw, S., Augustijn, H. E., Reitz, Z. L., Biermann, F., Alanjary, M., Fetter, A. et al. 2023. antiSMASH 7.0: new and improved predictions for detection, regulation, chemical structures and visualisation. Nucleic Acids Research 51:W46–W50.

Bolger, A. M., Lohse, M., and Usadel, B. 2014. Trimmomatic: a flexible trimmer for Illumina sequence data. Bioinformatics 30:2114–2120.

Calvo, A. M., Lohmar, J. M., Ibarra, B., and Satterlee, T. 2016. 18 Velvet Regulation of Fungal Development. In Growth, Differentiation and Sexuality, edited by Jürgen Wendland, 475–497. Cham: Springer International Publishing.

Casey, T., Solomon, P. S., Bringans, S., Tan, K. C., Oliver, R. P., and Lipscombe, R. 2010. Quantitative proteomic analysis of G-protein signalling in Stagonospora nodorum using isobaric tags for relative and absolute quantification. Proteomics 10:38–47.

Charoensawan, V., Wilson, D., and Teichmann, S. A. 2010. Genomic repertoires of DNA-binding transcription factors across the tree of life. Nucleic Acids Res 38:7364–77.

Chooi, Y.-H., Muria-Gonzalez Mariano, J., Mead Oliver, L., and Solomon Peter, S. 2015. SnPKS19 Encodes the polyketide synthase for alternariol mycotoxin biosynthesis in the wheat pathogen Parastagonospora nodorum. Applied and Environmental Microbiology 81:5309–5317.

Chooi, Y.-H., Zhang, G., Hu, J., Muria-Gonzalez, M. J., Tran, P. N., Pettitt, A., Maier, A. G., Barrow, R. A., and Solomon, P. S. 2017. Functional genomics-guided discovery of a light-activated phytotoxin in the wheat pathogen Parastagonospora nodorum via pathway activation. Environmental Microbiology 19:1975–1986.

Chooi, Y. H., Muria-Gonzalez, M. J., and Solomon, P. S. 2014. A genome-wide survey of the secondary metabolite biosynthesis genes in the wheat pathogen Parastagonospora nodorum. Mycology 5:192–206.

Consortium, T. U. 2022. UniProt: the Universal Protein Knowledgebase in 2023. Nucleic Acids Research 51:D523–D531.

Dobin, A., Davis, C. A., Schlesinger, F., Drenkow, J., Zaleski, C., Jha, S., Batut, P., Chaisson, M., and Gingeras, T. R. 2013. STAR: ultrafast universal RNA-Seq aligner. Bioinformatics 29:15–21.

Downie, R. C., Lin, M., Corsi, B., Ficke, A., Lillemo, M., Oliver, R. P., Phan, H. T. T., Tan, K.-C., and Cockram, J. 2020. Septoria nodorum blotch of wheat: disease management and resistance breeding in the face of shifting disease dynamics and a changing environment. Phytopathology® 111:906–920.

Duran, R. M., Cary, J. W., and Calvo, A. M. 2007. Production of cyclopiazonic acid, aflatrem, and aflatoxin by Aspergillus flavus is regulated by veA, a gene necessary for sclerotial formation. Applied Microbiology and Biotechnology 73:1158–1168.

Engler, C., Kandzia, R., and Marillonnet, S. 2008. A one pot, one step, precision cloning method with high throughput capability. PLOS ONE 3:e3647.

Eom, T. J., Moon, H., Yu, J. H., and Park, H. S. 2018. Characterization of the velvet regulators in Aspergillus flavus. J Microbiol 56:893–901.

Faris, J. D., and Friesen, T. L. 2020. Plant genes hijacked by necrotrophic fungal pathogens. Current Opinion in Plant Biology 56:74–80.

Friesen, T. L., Stukenbrock, E. H., Liu, Z., Meinhardt, S., Ling, H., Faris, J. D., Rasmussen, J. B., Solomon, P. S., McDonald, B. A., and Oliver, R. P. 2006. Emergence of a new disease as a result of interspecific virulence gene transfer. Nature Genetics 38:953–956.

Friesen, T. L., Zhang, Z., Solomon, P. S., Oliver, R. P., and Faris, J. D. 2008. Characterization of the interaction of a novel Stagonospora nodorum host-selective toxin with a wheat susceptibility gene. Plant Physiol 146:682–93.

Fulton, T. R., Ibrahim, N., Losada, M. C., Grzegorski, D., and Tkacz, J. S. 1999. A melanin polyketide synthase (PKS) gene from Nodulisporium sp. that shows homology to the pks1 gene of Colletotrichum lagenarium. Molecular and General Genetics MGG 262:714–720.

Graziani, S., Vasnier, C., and Daboussi, M. J. 2004. Novel polyketide synthase from Nectria haematococca. Appl Environ Microbiol 70:2984–8.

Harishchandra, D. L., Zhang, W., Li, X., Chethana, K. W. T., Hyde, K. D., Brooks, S., Yan, J., and Peng, J. 2020. A LysM domain-containing protein LtLysM1 is important for vegetative growth and pathogenesis in woody plant pathogen Lasiodiplodia theobromae. Plant Pathol J 36:323–334.

Hilgarth, R. S., and Lanigan, T. M. 2020. Optimization of overlap extension PCR for efficient transgene construction. MethodsX 7:100759.

Ipcho, S. V., Hane, J. K., Antoni, E. A., Ahren, D., Henrissat, B., Friesen, T. L., Solomon, P. S., and Oliver, R. P. 2012. Transcriptome analysis of Stagonospora nodorum: gene models, effectors, metabolism and pantothenate dispensability. Mol Plant Pathol 13:531–45.

IpCho, S. V., Tan, K. C., Koh, G., Gummer, J., Oliver, R. P., Trengove, R. D., and Solomon, P. S. 2010. The transcription factor StuA regulates central carbon metabolism, mycotoxin production, and effector gene expression in the wheat pathogen Stagonospora nodorum. Eukaryot Cell 9:1100–8.

John, E., Jacques, S., Phan, H. T. T., Liu, L., Pereira, D., Croll, D., Singh, K. B., Oliver, R. P., and Tan, K.-C. 2022. Variability in an effector gene promoter of a necrotrophic fungal pathogen dictates epistasis and effector-triggered susceptibility in wheat. PLOS Pathogens 18:e1010149.

John, E., Singh, K. B., Oliver, R. P., and Tan, K.-C. 2021. Transcription factor control of virulence in phytopathogenic fungi. Molecular Plant Pathology 22:858–881.

Jones, D. A. B., John, E., Rybak, K., Phan, H. T. T., Singh, K. B., Lin, S.-Y., Solomon, P. S., Oliver, R. P., and Tan, K.-C. 2019. A specific fungal transcription factor controls effector gene expression and orchestrates the establishment of the necrotrophic pathogen lifestyle on wheat. Scientific Reports 9:15884.

Jones, D. A. B., Rybak, K., Bertazzoni, S., Tan, K.-C., Phan, H. T. T., and Hane, J. K. 2021. Pathogenicity effector candidates and accessory genome revealed by pan-genomic analysis of Parastagonospora nodorum. bioRxiv:2021.09.01.458590.

Käfer, E. 1965. Origins of translocations in Aspergillus nidulans. Genetics 52:217–32.

Kettles, G. J., Bayon, C., Sparks, C. A., Canning, G., Kanyuka, K., and Rudd, J. J. 2018. Characterization of an antimicrobial and phytotoxic ribonuclease secreted by the fungal wheat pathogen Zymoseptoria tritici. New Phytol 217:320–331.

Kim, H.-J., Han, J.-H., Kim, K. S., and Lee, Y.-H. 2014. Comparative functional analysis of the velvet gene family reveals unique roles in fungal development and pathogenicity in Magnaporthe oryzae. Fungal Genetics and Biology 66:33–43.

Kombrink, A., Rovenich, H., Shi-Kunne, X., Rojas-Padilla, E., van den Berg, G. C., Domazakis, E., de Jonge, R. et al. 2017. Verticillium dahliae LysM effectors differentially contribute to virulence on plant hosts. Mol Plant Pathol 18:596–608.

Kombrink, A., and Thomma, B. P. 2013. LysM effectors: secreted proteins supporting fungal life. PLoS Pathog 9:e1003769.

Li, H., Wei, H., Hu, J., Lacey, E., Sobolev, A. N., Stubbs, K. A., Solomon, P. S., and Chooi, Y.-H. 2020. Genomics-driven discovery of phytotoxic cytochalasans involved in the virulence of the wheat pathogen Parastagonospora nodorum. ACS Chemical Biology 15:226–233.

Liao, Y., Smyth, G. K., and Shi, W. 2014. featureCounts: an efficient general purpose program for assigning sequence reads to genomic features. Bioinformatics 30:923–30.

Lin, S. Y., Chooi, Y. H., and Solomon, P. S. 2018. The global regulator of pathogenesis PnCon7 positively regulates Tox3 effector gene expression through direct interaction in the wheat pathogen Parastagonospora nodorum. Mol Microbiol.

Liu, Z. H., Faris, J. D., Meinhardt, S. W., Ali, S., Rasmussen, J. B., and Friesen, T. L. 2004. Genetic and physical mapping of a gene conditioning sensitivity in wheat to a partially purified host-selective toxin produced by Stagonospora nodorum. Phytopathology 94:1056–60.

Love, M. I., Huber, W., and Anders, S. 2014. Moderated estimation of fold change and dispersion for RNA-Seq data with DESeq2. Genome Biology 15:550.

Moon, H., Lee, M.-K., Bok, I., Bok, J. W., Keller, N. P., and Yu, J.-H. 2023. Unraveling the gene regulatory networks of the global regulators VeA and LaeA in Aspergillus nidulans. Microbiology Spectrum 11:e00166–23.

Morishita, Y., Aoki, Y., Ito, M., Hagiwara, D., Torimaru, K., Morita, D., Kuroda, T. et al. 2020. Genome mining-based discovery of fungal macrolides modified by glycosylphosphatidylinositol (GPI)– ethanolamine phosphate transferase homologues. Organic Letters 22:5876–5879.

Müller, N., Leroch, M., Schumacher, J., Zimmer, D., Könnel, A., Klug, K., Leisen, T. et al. 2018. Investigations on VELVET regulatory mutants confirm the role of host tissue acidification and secretion of proteins in the pathogenesis of Botrytis cinerea. New Phytologist 219:1062–1074.

Myung, K., Li, S., Butchko, R. A. E., Busman, M., Proctor, R. H., Abbas, H. K., and Calvo, A. M. 2009. FvVE1 Regulates biosynthesis of the mycotoxins Fumonisins and Fusarins in Fusarium verticillioides. Journal of Agricultural and Food Chemistry 57:5089–5094.

Oliver, R. P., Friesen, T. L., Faris, J. D., and Solomon, P. S. 2012. Stagonospora nodorum: from pathology to genomics and host resistance. Annu Rev Phytopathol 50:23–43.

Ostry, V. 2008. Alternaria mycotoxins: an overview of chemical characterization, producers, toxicity, analysis and occurrence in foodstuffs. World Mycotoxin Journal 1:175–188.

Park, P. J. 2009. ChIP–Seq: advantages and challenges of a maturing technology. Nature Reviews Genetics 10:669–680.

Paysan-Lafosse, T., Blum, M., Chuguransky, S., Grego, T., Pinto, B. L., Salazar, Gustavo A., Bileschi, Maxwell L. et al. 2022. InterPro in 2022. Nucleic Acids Research 51:D418–D427.

Piras, M., Patruno, I., Nikolakopoulou, C., Willment, J. A., Sloan, N. L., Zanato, C., Brown, G. D., and Zanda, M. 2021. Synthesis of the fungal metabolite YWA1 and related constructs as tools to study MelLec-mediated immune response to Aspergillus infections. The Journal of Organic Chemistry 86:6044–6055.

Quaedvlieg, W., Verkley, G. J. M., Shin, H. D., Barreto, R. W., Alfenas, A. C., Swart, W. J., Groenewald, J. Z., and Crous, P. W. 2013. Sizing up Septoria. Studies in Mycology 75:307–390.

Quevillon, E., Silventoinen, V., Pillai, S., Harte, N., Mulder, N., Apweiler, R., and Lopez, R. 2005. InterProScan: protein domains identifier. Nucleic Acids Res 33:W116–20.

Richards, J. K., Kariyawasam, G. K., Seneviratne, S., Wyatt, N. A., Xu, S. S., Liu, Z., Faris, J. D., and Friesen, T. L. 2022. A triple threat: the Parastagonospora nodorum SnTox267 effector exploits three distinct host genetic factors to cause disease in wheat. New Phytologist 233:427–442.

Rybak, K., See, P. T., Phan, H. T., Syme, R. A., Moffat, C. S., Oliver, R. P., and Tan, K. C. 2017. A functionally conserved Zn(2) Cys(6) binuclear cluster transcription factor class regulates necrotrophic effector gene expression and host-specific virulence of two major Pleosporales fungal pathogens of wheat. Mol Plant Pathol 18:420–434.

Savary, S., Willocquet, L., Pethybridge, S. J., Esker, P., McRoberts, N., and Nelson, A. 2019. The global burden of pathogens and pests on major food crops. Nature Ecology & Evolution 3:430–439.

Schumacher, J., Pradier, J.-M., Simon, A., Traeger, S., Moraga, J., Collado, I. G., Viaud, M., and Tudzynski, B. 2012. Natural variation in the VELVET gene bcvel1 affects virulence and light-dependent differentiation in Botrytis cinerea. PLOS ONE 7:e47840.

Sharpee, W., Oh, Y., Yi, M., Franck, W., Eyre, A., Okagaki, L. H., Valent, B., and Dean, R. A. 2017. Identification and characterization of suppressors of plant cell death (SPD) effectors from Magnaporthe oryzae. Mol Plant Pathol 18:850–863.

Solomon, P. S., Ipcho, S. V. S., Hane, J. K., Tan, K. C., and Oliver, R. P. 2008. A quantitative PCR approach to determine gene copy number. Fungal Genetics Reports 55:5–8.

Solomon, P. S., Lee, R. C., Wilson, T. J., and Oliver, R. P. 2004. Pathogenicity of Stagonospora nodorum requires malate synthase. Mol Microbiol 53:1065–73.

Solomon, P. S., Lowe, R. G. T., Tan, K.-C., Waters, O. D. C., and Oliver, R. P. 2006. Stagonospora nodorum: cause of stagonospora nodorum blotch of wheat. Molecular Plant Pathology 7:147–156.

Tamura, K., Stecher, G., and Kumar, S. 2021. MEGA11: Molecular Evolutionary Genetics Analysis version 11. Molecular Biology and Evolution 38:3022–3027.

Tan, K.-C., and Oliver, R. P. 2017. Regulation of proteinaceous effector expression in phytopathogenic fungi. PLOS Pathogens 13:e1006241.

Tan, K.-C., Trengove, R. D., Maker, G. L., Oliver, R. P., and Solomon, P. S. 2009. Metabolite profiling identifies the mycotoxin alternariol in the pathogen Stagonospora nodorum. Metabolomics 5:330–335.

Tan, K. C., Phan, H. T., Rybak, K., John, E., Chooi, Y. H., Solomon, P. S., and Oliver, R. P. 2015. Functional redundancy of necrotrophic effectors - consequences for exploitation for breeding. Front Plant Sci 6:501.

Wang, L., Wang, M., Jiao, J., and Liu, H. 2022. Roles of AaVeA on mycotoxin production via light in Alternaria alternata. Frontiers in Microbiology 13.

Wenderoth, M., Garganese, F., Schmidt-Heydt, M., Soukup, S. T., Ippolito, A., Sanzani, S. M., and Fischer, R. 2019. Alternariol as virulence and colonization factor of Alternaria alternata during plant infection. Molecular Microbiology 112:131–146.

Yee, D. A., Niwa, K., Perlatti, B., Chen, M., Li, Y., and Tang, Y. 2023. Genome mining for unknown-unknown natural products. Nat Chem Biol 19:633–640.

Young, M. D., Wakefield, M. J., Smyth, G. K., and Oshlack, A. 2010. Gene ontology analysis for RNA-Seq: accounting for selection bias. Genome Biology 11:R14.

